# Opposing actions of *cis-* and *trans-*ligands at a single GPCR link osmotic state to habitat choice

**DOI:** 10.64898/2026.06.29.735401

**Authors:** Yuma Tsukasa, Takumi S. Ohbayashi, Misato Kurio, Hiroshi Kohsaka, Kaho Maeta, Etsu Matsutani, Akimitsu Tsumadori, Kai Li, Risa Matsui, Jun Suzuki, Akinao Nose, Tadashi Uemura, Tadao Usui

## Abstract

How internal physiological states are translated into altered behavioral responses to the same sensory cue remains poorly understood. Here we show that the glycosylphosphatidylinositol (GPI)-anchored Ly6/uPAR protein Belly roll (Bero) acts as a *cis*-ligand for the neuropeptide receptor Allatostatin C receptor 2 (AstC-R2) in Drosophila larvae. Bero–AstC-R2 signaling establishes a phospholipase Cβ-dependent high-Ca²⁺ state in abdominal leucokinin-producing (ABLK) neurons that suppresses nociceptive sensory transmission. Desiccation raises hemolymph osmolality, which activates AstC-producing neurons; the secreted AstC peptide then opposes Bero–AstC-R2 signaling, lowers ABLK Ca²⁺ activity, and unlocks avoidance of dry substrates. As a consequence, dehydrated larvae bias their pupation sites toward humid environments, mitigating desiccation risk during the immobile pupal stage. Thus, a membrane-tethered *cis*- and a secreted *trans*-ligand can act in opposition at a single GPCR, providing a non-canonical mechanism by which organismal physiological state is encoded in receptor signaling and converted into an adaptive behavioral choice.

## INTRODUCTION

Animals continually adjust their behavior to meet their physiological needs. Internal physiological states, such as hunger and thirst, alter how the same sensory stimuli are interpreted, enabling animals to select adaptive behaviors and maintain homeostasis in changing environments.^1,2^ Dehydration provides a particularly stringent example of this state-dependent regulation: even small changes in body-fluid osmolality can drive thirst and water-seeking behavior, whereas larger perturbations can severely compromise physiological function.^3,4^ Changes in body-fluid osmolality alter neural activity and promote water-directed or humidity-seeking behaviors in both mammals^5–7^ and insects.^8–10^ However, how holometabolous insects cope with the risk of dehydration during the pupal stage—when they are unable to move in search of water—remains unclear.

Last-instar larvae of most holometabolous insects actively select sites for pupation. For example, *Drosophila melanogaster* larvae adjust their pupation sites in response to environmental cues such as ambient light, temperature, and humidity.^11–13^ However, whether internal physiological states such as dehydration influence pupation-site selection remains unclear,^14^ as previous studies have not distinguished the acute effects of environmental fluctuations from the chronic effects of internal states.^11^ We therefore hypothesized that *Drosophila* larvae use their dehydration state as a cue to predict potential desiccation stress during the upcoming pupal stage and mitigate this risk by adjusting their pupation sites in advance. Such behavioral flexibility is often mediated by neuromodulatory systems.^1,15–19^

One class of candidate neuromodulatory molecules is the lymphocyte antigen-6 (Ly6)/urokinase-type plasminogen activator receptor (uPAR) superfamily (LU superfamily) of glycosylphosphatidylinositol (GPI)-anchored proteins. Most members of the LU superfamily are anchored to the outer leaflet of the cell membrane via a GPI moiety.^20,21^ Through their extracellular three-finger domains, LU proteins interact in *cis* with ion channels and transmembrane receptors to modulate cellular excitability and signaling.^22–27^ Although several LU superfamily proteins have been reported to form complexes with G protein-coupled receptors (GPCRs),^28–32^ whether endogenous LU proteins can act as *cis*-ligands for GPCRs to modulate intracellular signaling and animal behavior remains unclear. We previously showed that the LU superfamily protein Belly roll (Bero) regulates the activity of abdominal leucokinin-producing (ABLK) neurons,^33,34^ which control nociceptive avoidance behaviors;^33,35–37^ however, the molecular target of Bero remains unknown.

Here, we investigated dehydration-dependent habitat navigation in *Drosophila* larvae and found that internal osmotic state shapes avoidance behavior through a non-canonical neuromodulatory mechanism. Desiccation increases hemolymph osmolality and enhances avoidance of dry substrates, thereby biasing pupation-site selection toward wetter environments. This behavioral switch involves ABLK neurons, in which Bero acts in *cis* with the GPCR Allatostatin C receptor 2 (AstC-R2). Under low-osmolality conditions, Bero–AstC-R2 signaling establishes a phospholipase Cβ (PLCβ)-dependent cellular state characterized by elevated Ca²⁺ activity that suppresses sensory transmission. Under high-osmolality conditions, secreted Allatostatin C (AstC) counteracts Bero–AstC-R2 signaling, reducing ABLK Ca²⁺ activity and facilitating dry-substrate avoidance. Our findings reveal a non-canonical neuromodulatory mechanism involving opposing actions of a membrane-tethered and a secreted ligand at a single GPCR, through which internal osmotic state reshapes pupation-site navigation behavior.

## RESULTS

### Desiccation experience alters larval pupation-site navigation

Desiccation is a major environmental stressor for small terrestrial animals, particularly insects with small body sizes and high surface area-to-volume ratios, which promote greater water loss relative to body mass.^38–43^ Although previous studies have shown that *Drosophila* larvae alter their pupation sites in response to light, humidity, or substrate water content, whether desiccation-induced changes in internal state influence larval navigation remains unclear.^11–13,44–47^

To test this possibility, we exposed larvae to desiccation stress for 1 h (<20% relative humidity [RH]; “desiccation phase”) and then transferred them to a humidified vertical food vial (>90% RH; “navigation phase”). After pupariation, the height of each pupa above the food surface was measured (“pupation height”; see Figures 1A and S1A). Because the lower portion of the plastic surface is wetter than the upper portion,^11^ the pupation site selected by each larva serves as a proxy for its preference for relatively dry versus wet substrates. Following exposure to desiccation stress, larvae selected lower regions of the vial wall as pupation sites during the navigation phase (Figure 1B; Video S1). This shift was not attributable to impaired locomotion, as desiccation-experienced larvae were able to climb to the top of agar-coated vial walls (Figures S1C and S1D). These findings indicate that prior desiccation experience alters pupation-site navigation in a substrate-dependent manner, suggesting that internal physiological state influences subsequent habitat selection

### Dehydration facilitates dry-substrate avoidance

DesWe next asked which larval behaviors underlie the selection of lower pupation sites following desiccation. We focused on head- and/or tail-raising (HTR) behavior,^48^ as wandering larvae frequently raised their heads and/or tails on dry plastic walls and occasionally fell from the surface (Figure S2A; Video S2). To characterize HTR behavior more precisely, we placed larvae on a horizontally mounted dry plastic plate and recorded their movements from the side (Figure 1C; Video S3). On dry surfaces, larvae attempted to crawl but often failed to move smoothly and occasionally raised their heads and/or tails (Figure 1D). These behavioral responses occurred not only on dry plastic surfaces but also on fruit surfaces, which represent natural food substrates for larvae (Figure S2B). Moreover, wild *Drosophila* larvae exhibited HTR behavior on dry surfaces, such as wood chips and fallen leaves, under field conditions (Figures S2C and S2D; Video S4). By contrast, we did not observe HTR behavior on wet surfaces, such as agar-coated plastic substrates. Together, these observations suggest that HTR behavior is associated with larval responses to dry substrates.

**Figure 1.**
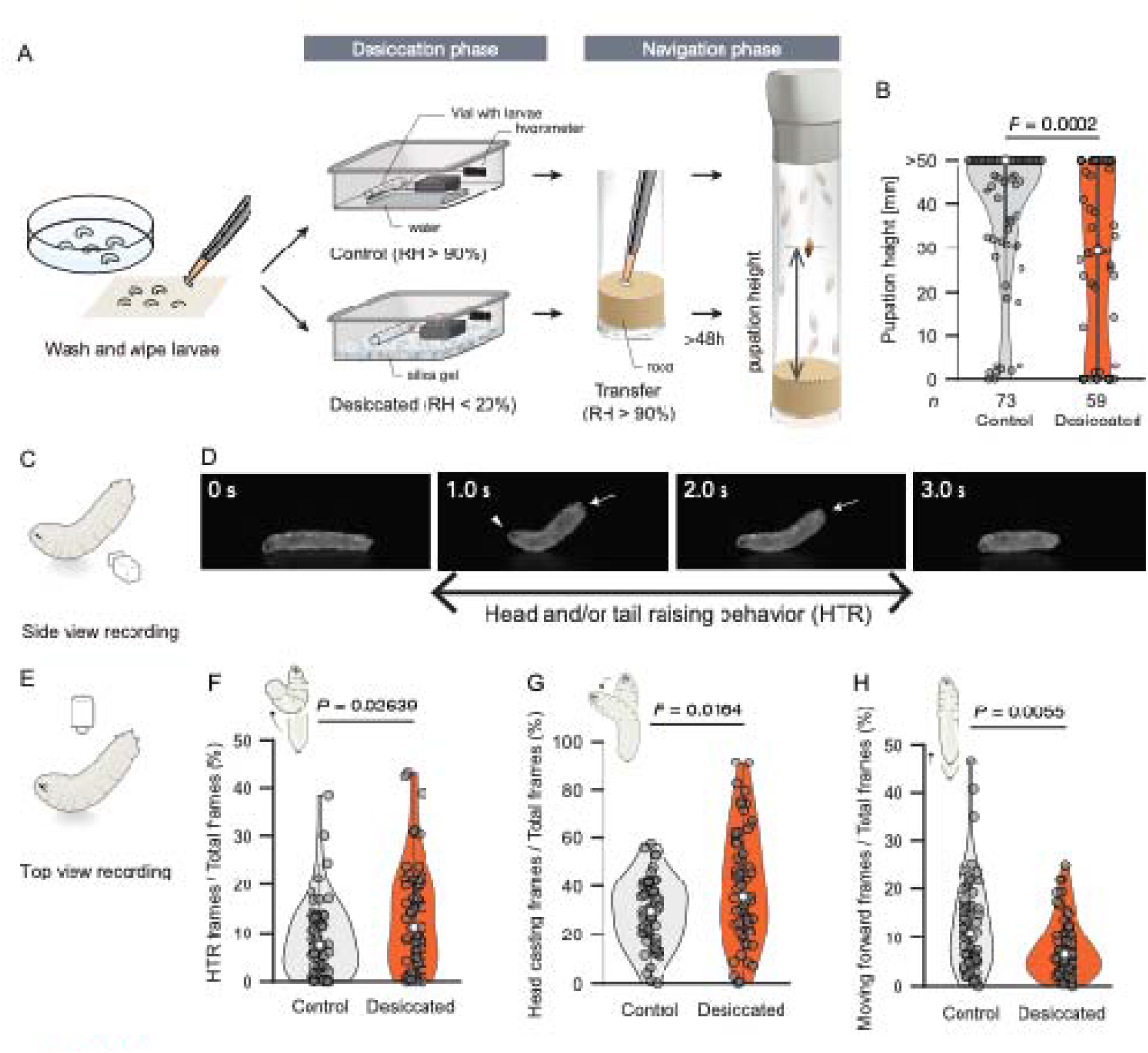
Dehydration promotes dry-substrate avoidance and modulates pupation-site navigation. (A) Schematic of the vertical pupation-site assay. During the desiccation phase, control larvae were exposed to >90% relative humidity (RH) for 1 h, whereas desiccation-experienced larvae were exposed to <20% RH. Larvae were then transferred to a humidified vertical food vial for the navigation phase. Illustrations were adapted and modified from Kurio et al.^64^ with permission from the publisher. (B) Quantification of pupation height in control and desiccation-experienced larvae. Wilcoxon rank-sum test. (C) Schematic of side-view recording on a dry plastic surface. (D) Representative frames from a side-view recording. Arrowhead, head raising; arrows, tail raising. (E) Schematic of top-view recording for behavioral classification. (F to H) Quantification of larval behavior on dry substrate surfaces during 1-min recordings: proportion of frames showing head- and/or tail-raising (HTR) behavior (F), head casting (G), and forward movement (H). Control, *n* = 55; desiccation-experienced, *n* = 57. Wilcoxon rank-sum test. See also Figures S1-S6.

To determine how prior desiccation experience influences larval behavior on dry substrates, including HTR behavior, we analyzed larval movements using top-view recordings (Figure 1E). We developed a machine learning-based classification system to reliably detect HTR behavior from video recordings at high throughput (Figure S3; Video S5). Using this system, we found that HTR behavior increased following desiccation experience (Figures 1F and S4A). Desiccation-experienced larvae also exhibited more head casting (Figure 1G), indicative of attempts to alter their trajectory,^49,50^ while spending less time moving forward (Figure 1H). Nonetheless, straight-line distance and average velocity did not differ between control and desiccation-experienced larvae (Figures S4B and S4C). These findings are consistent with the observation that prior desiccation does not impair general larval locomotor ability (Figures S1C and S1D).

**Figure 2.**
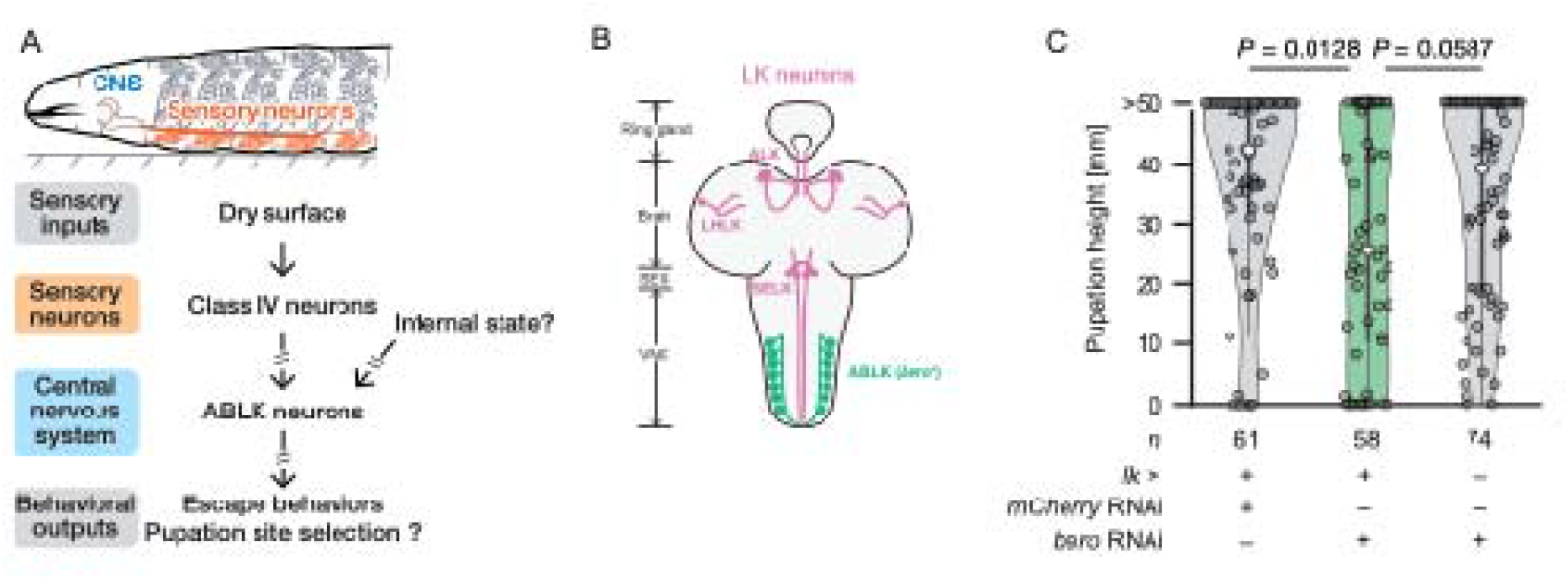
Bero in ABLK neurons modulates pupation-site navigation. (A) Schematic of larval Class IV sensory neurons (orange) and the central nervous system (CNS; blue) containing abdominal leucokinin-producing (ABLK) neurons. Proposed model in which dry-substrate sensory input from Class IV neurons and a hypothesized internal osmotic-state signal resulting from prior desiccation experience converge on ABLK neurons to control pupation-site selection. (B) Schematic of larval *lk^+^* neurons; image modified from Kurio et al.^33^ The larval CNS contains ABLK neurons, anterior LK (ALK) neurons, lateral horn LK (LHLK) neurons, and subesophageal LK (SELK) neurons. Among these *lk*^+^ neuronal populations, Bero expression is restricted to ABLK neurons. (C) Pupation height of control larvae (LK > *mCherry* RNAi or + > *bero* RNAi) and larvae with *bero* knocked down in LK neurons (LK > *bero* RNAi). Kruskal–Wallis test, followed by Dunn’s test. See also Figure S7.

We next asked how changes in short-timescale behavioral features, including HTR behavior, head casting, and forward movement, contribute to altered pupation-site selection following desiccation. To link these behaviors to pupation-site selection, we constructed an agent-based model in which simulated larvae exhibited different behavioral properties—HTR behavior, head casting, forward movement, and stopping—according to whether they were in contact with a wet or dry surface (Figure S5A). Simulated larvae were also programmed to wet the surface as they moved (Figure S5B).^11^ The model largely recapitulated the lower pupation sites observed in desiccation-experienced larvae and predicted increased pupation on wet surface regions (Figures S6A, SB, and SE; simulation results shown for *P*_falling_ = 1.0 × 10^−1.7^).

It was not possible to determine experimentally whether each larva selected a wet or dry surface for pupation because the wetness of the plastic wall could not be accurately assessed during the wandering stage. To address this uncertainty, we used a simulation-based estimation method to infer surface wetness. The predicted proportion of larvae selecting wet surfaces was higher among dehydrated larvae across the estimated range of *P*_falling_ (Figure S6C). Although the estimated wet surface area did not differ significantly at *P*_falling_ = 1.0 × 10^−1.7^, a larger proportion of desiccation-experienced larvae than control larvae selected wet surfaces as pupation sites (Figures S6D, F, and G).

To assess the contribution of HTR behavior to pupation-site selection, we introduced a hypothetical constraint into the simulation in which simulated individuals did not exhibit HTR behavior. Removing HTR behavior did not affect the median pupation height, regardless of desiccation experience (Figures S6H). However, it reduced the wet surface area on the wall, thereby allowing a larger proportion of dehydrated larvae to enter dry-surface regions (Figures S6I and J). Together, the agent-based model suggests that these behavioral components jointly shape pupation-site preferences in wandering larvae.

### Bero in ABLK neurons modulates pupation-site navigation

Dry surfaces are detected by nociceptive Class IV sensory neurons,^11^ which elaborate dendritic arbors beneath the body wall epidermis.^51–57^ However, the central neurons that transform dry-surface nociceptive input into avoidance behavior remain poorly defined. To investigate how internal physiological states modulate behavioral output *in vivo*, we focused on ABLK neurons, which receive Class IV-mediated sensory input and regulate several nociceptive escape behaviors (Figure 2A), including rolling.^33,35–37,58,59^ Moreover, ABLK neurons in adult flies have been reported to respond to desiccation stress.^60,61^ We therefore hypothesized that larval ABLK neurons integrate sensory input elicited by dry substrates with internal-state signals induced by prior desiccation exposure (Figure 2A). As previously reported, Belly roll (Bero), a member of the LU superfamily, modulates ABLK neuronal activity and suppresses rolling behavior (Figure 2B). ^33,34^ To determine whether Bero in ABLK neurons contributes to pupation-site navigation, we knocked down *bero* in these neurons. *bero* knockdown significantly reduced pupation height compared with that in control larvae, resulting in pupation heights comparable to those observed in desiccation-experienced larvae (Figures 2C and S7). These results indicate that Bero signaling in ABLK neurons contributes to the regulation of pupation-site navigation.

### ABLK neurons integrate hemolymph osmolality shifts and nociceptive input

**Figure 3.**
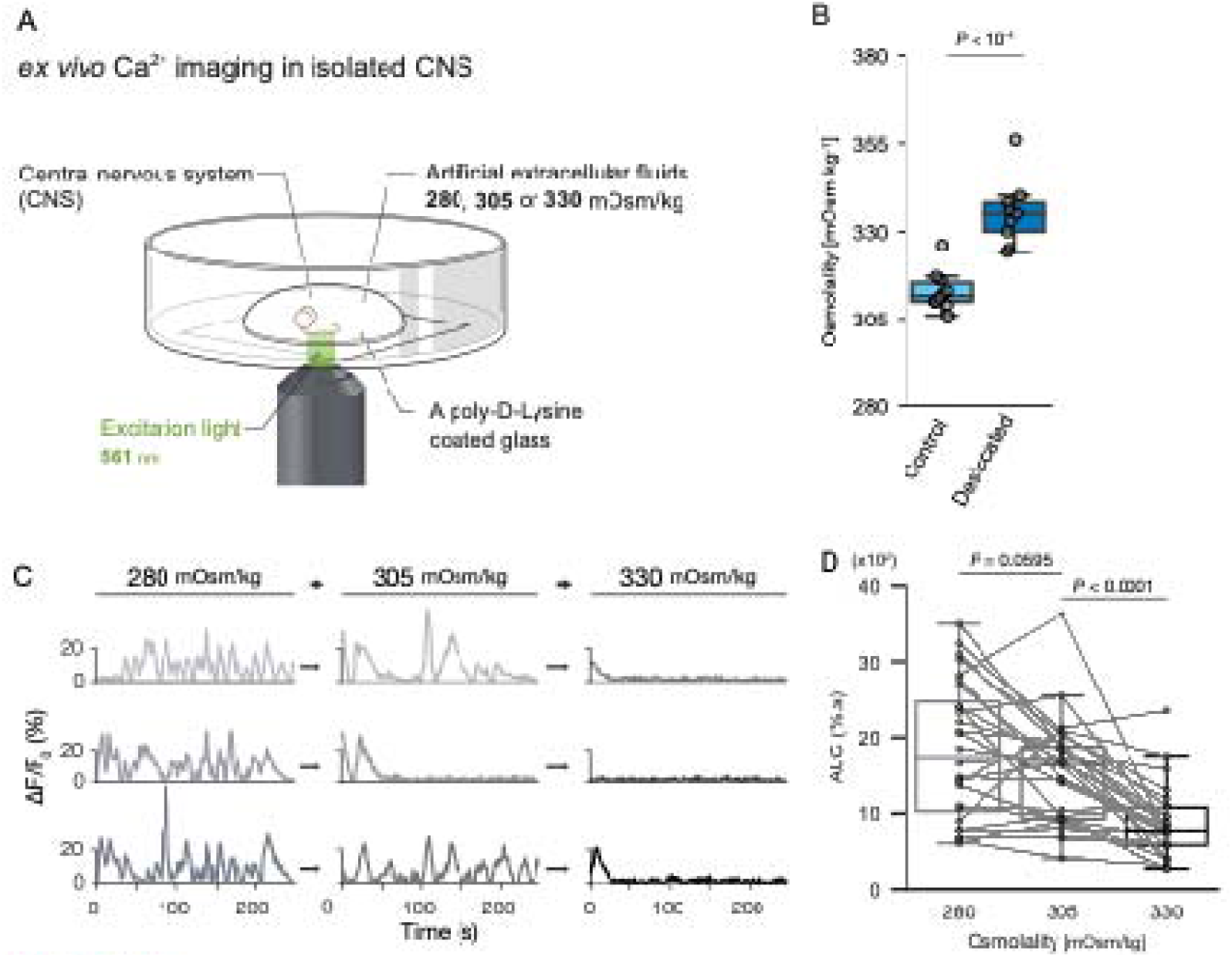
ABLK neurons are sensitive to changes in hemolymph osmolality. (A) Schematic of *ex vivo* Ca²⁺ imaging in isolated larval CNS preparations. (B) Hemolymph osmolality in control and desiccation-experienced larvae (control, *n* = 10; desiccation-experienced, *n* = 9). Wilcoxon rank-sum test. (C) Representative traces of spontaneous fluctuating Ca²⁺ activity (Δ *F*_Persistent_/*F*_0_) in ABLK neurons during sequential exposure to saline solutions with osmolalities of 280, 305, and 330 mOsm/kg. (D) Quantification of spontaneous Ca²⁺ activity, measured as the area under the curve (AUC), across osmolality conditions (*n* = 28 neurons from four larvae). Wilcoxon signed-rank test. See also Figure S8.

Desiccation stress increases body fluid osmolality in terrestrial animals.^62,63^ In adult *Drosophila*, interoceptive neurons in the subesophageal zone, lateral horn LK (LHLK) neurons, and subesophageal LK (SELK) neurons alter their activity in response to changes in hemolymph osmolality associated with thirst.^9,62^ We therefore investigated whether larval ABLK neurons are sensitive to changes in hemolymph osmolality. To address this question, we performed *ex vivo* Ca²⁺ imaging of isolated larval central nervous system (CNS) preparations immersed in artificial saline solutions of different osmolalities (Figure 3A). The hemolymph osmolality of wandering larvae has previously been estimated to be approximately 306.0 mOsm/kg.^64,65^ Here, we found that desiccation stress increased larval hemolymph osmolality by 23.7 mOsm/kg (Figure 3B). We therefore prepared artificial saline solutions with osmolalities of 280, 305, and 330 mOsm/kg by adjusting the concentrations of sucrose and trehalose, commonly used osmolytes;^9^ these solutions are hereafter referred to as S-280, S-305, and S-330, respectively. In S-280, ABLK neurons exhibited spontaneous Ca²⁺ activity (Figures 3C and 3D). When the recording saline was sequentially changed from S-280 to S-305 and then to S-330, spontaneous Ca²⁺ activity significantly decreased in the majority of ABLK neurons (Figures 3C and 3D). Furthermore, spontaneous Ca²⁺ activity was attenuated when osmolality was increased using l-glucose, a non-metabolizable osmolyte, indicating that the responses reflected increased osmolality rather than sugar metabolism (Figures S8A and S8B).

As previously reported, larval ABLK neurons exhibit two distinct types of activity: (1) spontaneous fluctuations in Ca²⁺ levels^33,37^ and (2) Class IV neuron-mediated sensory responses (see also Figure S10A).^33,36^ Because spontaneous Ca²⁺ activity was sensitive to osmolality, we next investigated whether evoked sensory responses in the same neurons were also modulated by osmolality. We monitored ABLK activity in semi-intact preparations using Ca²⁺ imaging (Figure 4A) and optogenetically activated Class IV neurons to mimic sensory input triggered by a dry substrate. Consistent with the results obtained in the isolated CNS, spontaneous ABLK activity changed in response to shifts in osmolality (control in Figure 4B; Video S6). In S-280, optogenetic activation of Class IV neurons did not elicit a detectable evoked response in ABLK neurons (control in Figures 4C and 4D). By contrast, ABLK neurons exhibited robust evoked responses in S-330 (Figures 4C and 4D). These reciprocal changes in spontaneous and evoked Ca²⁺ activity suggest that extracellular osmolality modulates the responsiveness of ABLK neurons to nociceptive sensory input.

**Figure 4.**
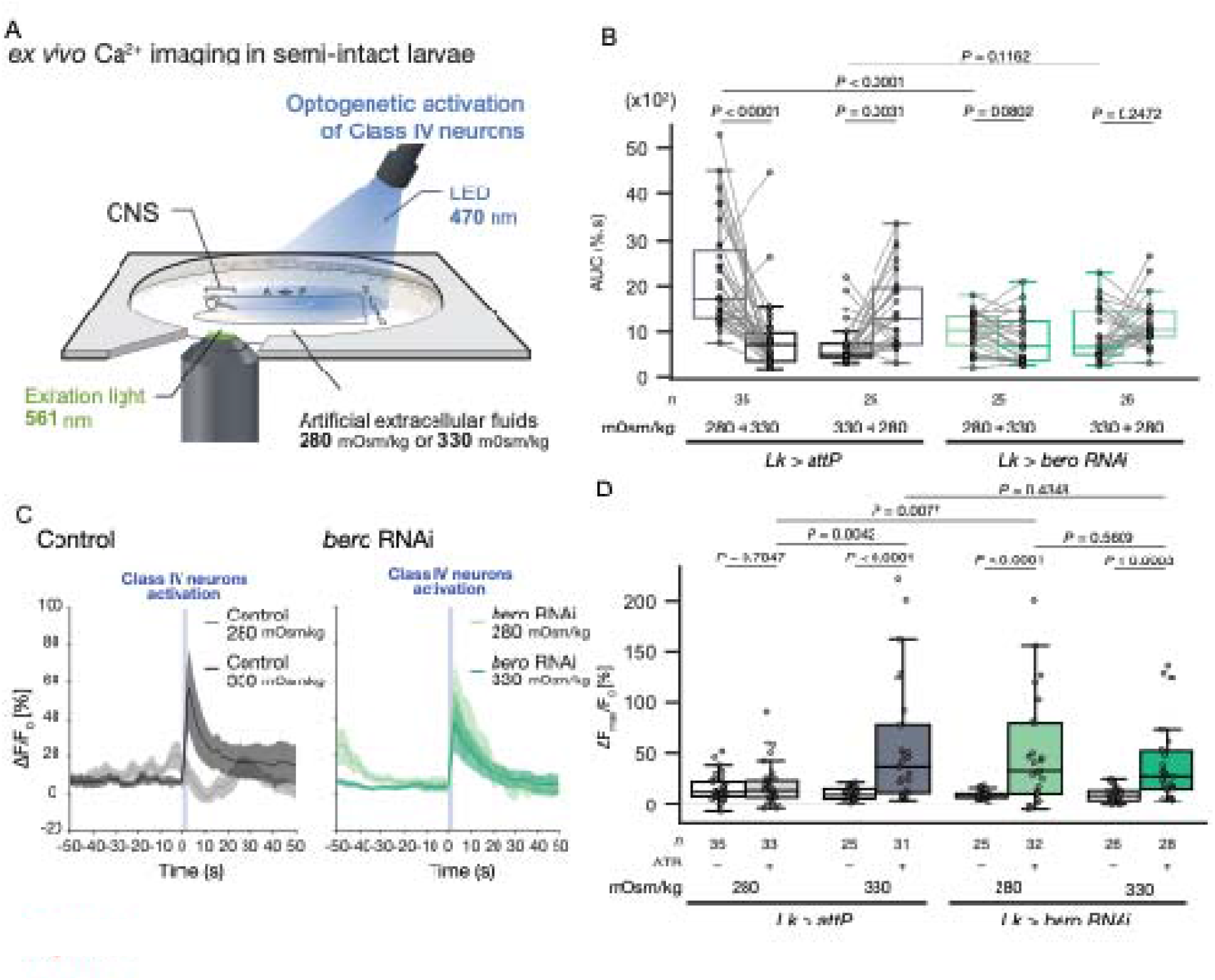
ABLK neurons gate nociceptive responses according to osmolality. (A) Schematic of Ca²⁺ imaging in semi-intact preparations with optogenetic activation of Class IV neurons. (B) Quantification of spontaneous Ca²⁺ activity, measured as AUC, during shifts between 280 and 330 mOsm/kg in control and *bero* knockdown larvae. Wilcoxon rank-sum test for unpaired comparisons and Wilcoxon signed-rank test for paired comparisons. (C) Representative evoked Ca²⁺ responses following optogenetic activation of Class IV neurons in control and *bero* knockdown larvae at 280 and 330 mOsm/kg. (D) Quantification of the maximum evoked Ca²⁺ response under the conditions shown in (G). Kruskal– Wallis test, followed by Dunn’s test. See also Figure S9 and S10.

Fluid osmolality can broadly influence neural activity and contribute to the regulation of internal states in mammals.^66,67^ We therefore asked whether the osmolality-dependent modulation observed in ABLK neurons was also present in other neurons within the larval nociceptive circuit. We examined A08n and SELK neurons. A08n neurons are second-order neurons within the nociceptive circuit and are located upstream of ABLK neurons,^35,36,68^ whereas SELK neurons represent another class of sensory interneurons.^9,33,36^ Unlike ABLK neurons, neither A08n nor SELK neurons exhibited detectable spontaneous activity under these experimental conditions. Moreover, their Class IV neurons-mediated sensory responses did not change across extracellular osmolalities tested (Figure S9). Collectively, these findings suggest that ABLK neurons specifically gate nociceptive sensory transmission through an osmolality-dependent mechanism.

**Figure 5.**
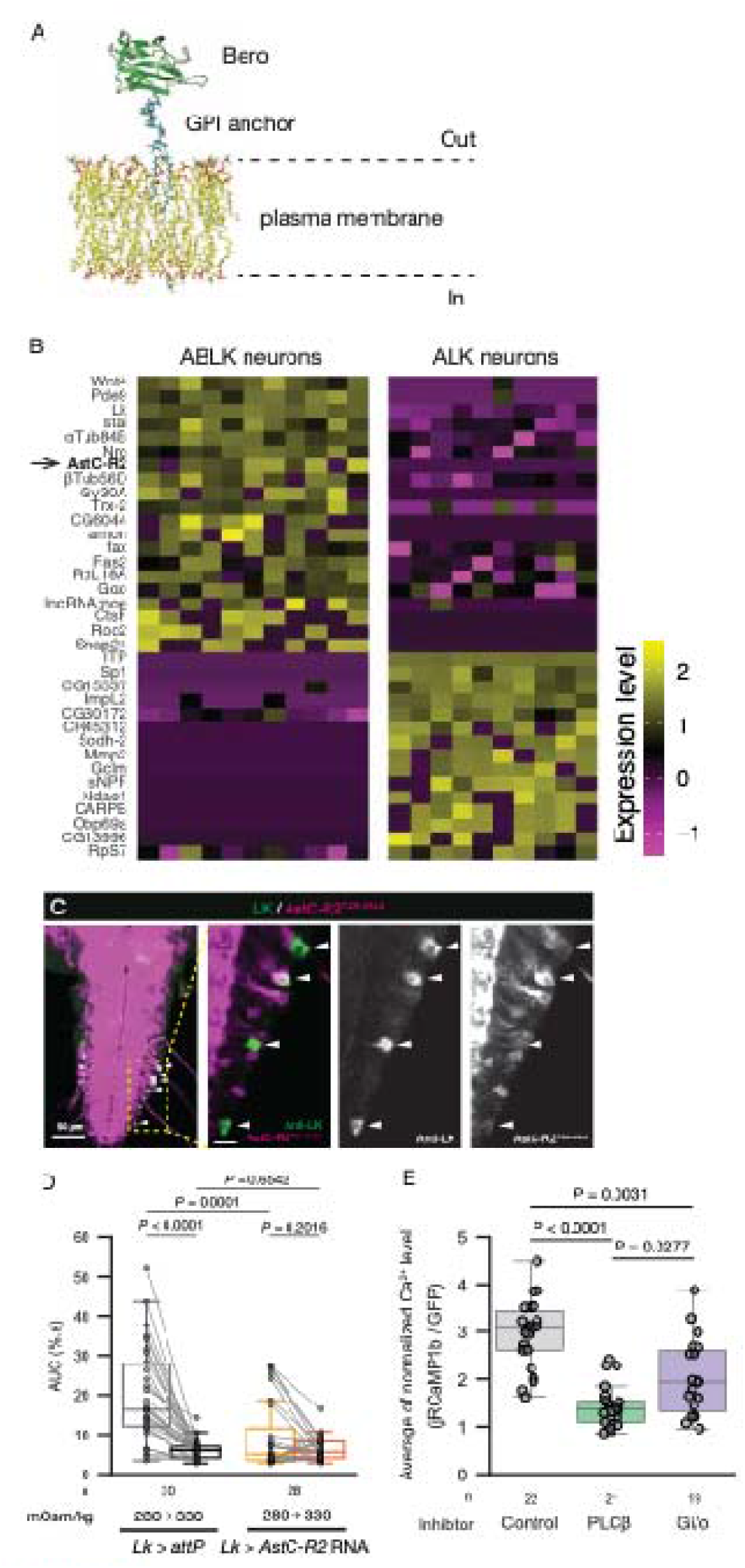
Bero and AstC-R2 establish a high-Ca²⁺ activity state through PLCβ-dependent signaling in ABLK neurons. (A) Predicted three-dimensional structure of Bero generated using AlphaFold2 and modeled in the plasma membrane using CHARMM-GUI. (B) Heat map showing the expression of marker genes and candidate transmembrane receptors in ABLK and ALK neurons from a publicly available dataset (GSE135810). (C) Confocal images showing *AstC-R2-T2A-GAL4*-labeled cells (magenta) and ABLK neurons (green). Scale bars, 50 μm and 10 μm. (D) Quantification of spontaneous Ca²⁺ activity, measured as AUC, in control and *AstC-R2* knockdown ABLK neurons at 280 and 330 mOsm/kg. Control*, n* = 30 neurons from three larvae; *AstC-R2* knockdown*, n* = 28 neurons from four larvae. Wilcoxon rank-sum test for unpaired comparisons and Wilcoxon signed-rank test for paired comparisons. (E) Quantification of normalized intracellular Ca²⁺ levels after treatment with the PLCβ inhibitor U73122 or pertussis toxin (PTX), an inhibitor of Gαi/o signaling, in Ca²⁺-free S-280. Kruskal–Wallis test, followed by Dunn’s test. See also Figure S11.

### Bero and AstC-R2 establish a high-Ca²⁺ activity state associated with low osmolality

We previously found that *bero* knockdown in ABLK neurons not only abolished spontaneous activity but also enhanced sensory responses in S-305,^33^ resembling the activity pattern of control neurons in S-330 (Figures 4B and 4D). We therefore investigated whether osmolality-dependent changes in ABLK neuronal activity require Bero. In S-280, *bero* knockdown abolished spontaneous activity (Figures 4B and S10A; Video S7) and significantly enhanced sensory responses (Figures 4C, 4D, and S10C). In S-330, however, *bero* knockdown did not further enhance sensory responses (Figures 4C, 4D, and S10C). These findings suggest that Bero is required to establish a high-Ca²⁺ activity state under low-osmolality conditions (hereafter, the “low-osmolality Ca²⁺ activity state”; see Figure S10) that suppresses evoked sensory responses. Notably, the low-Ca²⁺, high-osmolality-like state observed in ABLK neurons following *bero* knockdown was consistent with the enhanced dry-substrate avoidance reflected by lower pupation sites (Figures 2C and S7).

Because Bero lacks a cytoplasmic domain (Figure 5A), we reasoned that it may function through transmembrane proteins, such as ion channels or GPCRs.^33^ To identify candidate functional interactors of Bero in ABLK neurons, we searched single-cell RNA sequencing (scRNA-seq) datasets from the larval CNS^69,70^ for genes encoding transmembrane proteins expressed in ABLK neurons. This analysis identified *AstC-R2*, which encodes the neuropeptide GPCR AstC-R2, as a candidate Bero interactor (Figure 5B). An independent scRNA-seq dataset also supported *AstC-R2* expression in ABLK neurons.^71^ To further examine its expression pattern, we used an *AstC-R2-T2A-GAL4* reporter^72^ and found that reporter activity largely overlapped with ABLK neurons (Figure 5C). By contrast, neither SELK nor LHLK neurons were labeled (Figure S11A).

To determine whether AstC-R2 is required for the low-osmolality Ca²⁺ activity state in ABLK neurons, we knocked down *AstC-R2* in these neurons. *AstC-R2* knockdown nearly abolished spontaneous Ca²⁺ activity in S-280, consistent with loss of the low-osmolality Ca²⁺ activity state (Figures 5D and S11B). ABLK neurons have also been reported to be labeled by *5-HT1B-GAL4.*^37^ Unlike *AstC-R2* knockdown, however, *5-HT1B* knockdown did not abolish spontaneous activity (Figures S11C and S11D). In addition, a T2A-GAL4 reporter for *AstC-R1*, a paralog of *AstC-R2*, labeled ABLK neurons (Figure S11E), but *AstC-R1* knockdown did not alter the osmolality-dependent activity of ABLK neurons (Figures S11F and S11G). Together, these findings indicate that Bero and AstC-R2 are specifically required for the low-osmolality Ca²⁺ activity state in ABLK neurons.

**Figure 6.**
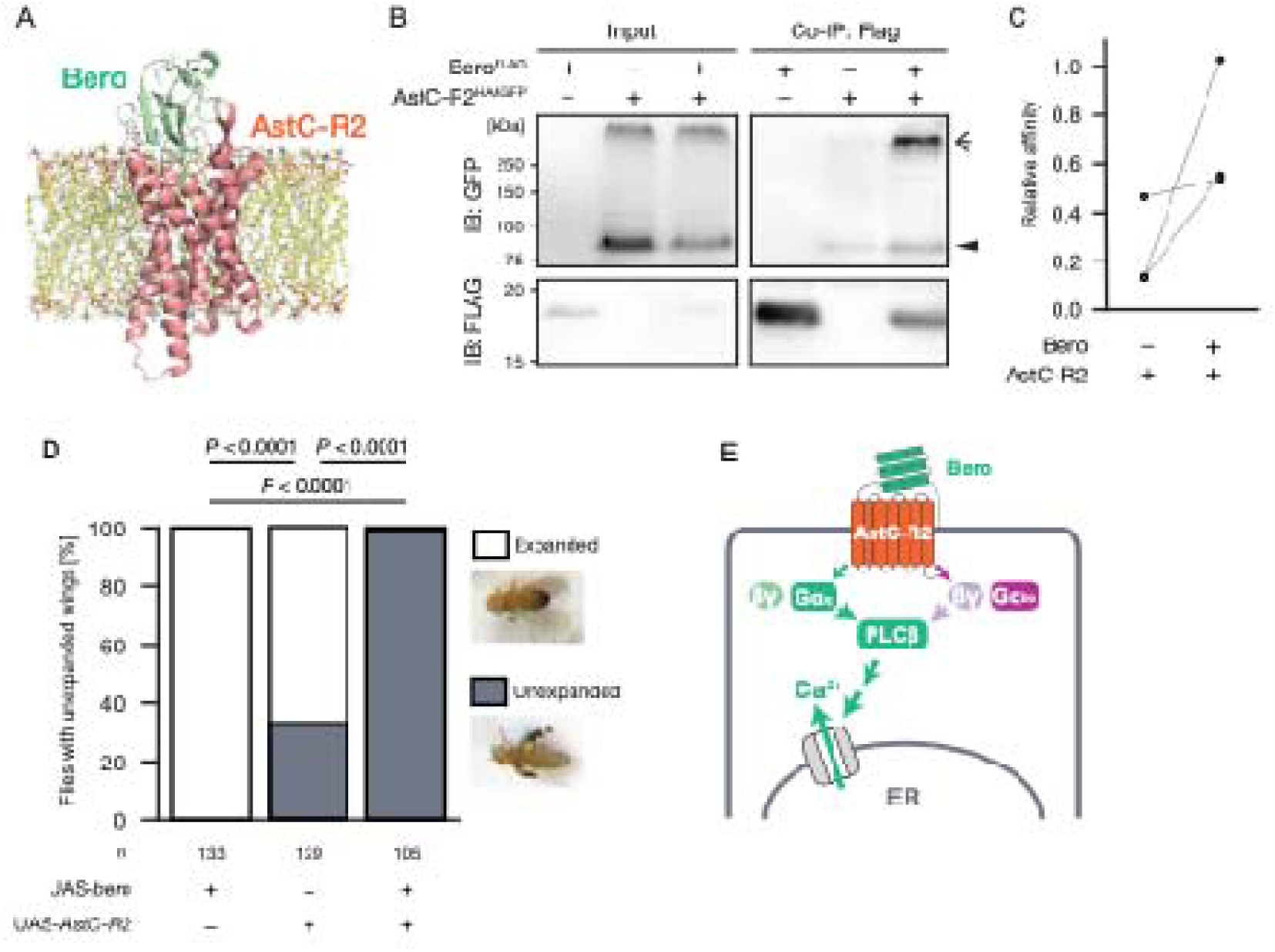
Bero and AstC-R2 form a functional *cis*-acting signaling module. (A) Predicted Bero–AstC-R2 complex structure generated using AlphaFold2-Multimer and modeled in a membrane environment using CHARMM-GUI. (B) Co-immunoprecipitation of Bero^FLAG^ with AstC-R2^HA&GFP^ in *Drosophila* S2 cells. Immunoblotting was performed using anti-GFP and anti-FLAG antibodies. The triangle indicates putative monomeric AstC-R2, and the arrow indicates a high-molecular-weight AstC-R2 species. (C) Quantification of the relative Bero–AstC-R2 association. (D) Proportion of adult flies with unexpanded wings following ectopic expression of Bero, AstC-R2, or both in type III neuroendocrine cells. (E) Proposed model in which membrane-tethered Bero modulates AstC-R2 signaling in *cis* to promote PLCβ-dependent Ca²⁺ activity. See also Figure S12–S5.

Although insect AstC receptors and mammalian somatostatin receptors (SSTRs) are generally associated with Gαi/o signaling,^73–77^ the Pred-Couple 2 algorithm^78^ predicted Gαq/11 as the most likely coupling partner for *Drosophila* AstC-R2 normalized posterior probability score (NPPS), 0.92), with Gαi/o as the second-ranked partner (NPPS, 0.11). Consistent with this prediction, pharmacological inhibition of PLCβ reduced Ca²⁺ activity in ABLK neurons, indicating that PLCβ is required for the low-osmolality Ca²⁺ activity state (Figure 5E). Pertussis toxin (PTX) also reduced Ca²⁺ activity, suggesting that Gαi/o and/or Gβγ signaling may contribute to this PLCβ-dependent activity.^79^ Together, these results indicate that the high-Ca²⁺ activity state of ABLK neurons requires PLCβ-dependent signaling, with possible contributions from both Gαq/11 and Gαi/o-associated pathways.

### Bero elicits AstC-R2 signaling through a cis-acting mechanism

Because both Bero and AstC-R2 are required for the low-osmolality Ca²⁺ activity state, we investigated whether the two proteins could form a molecular complex on the same cell membrane. We first used AlphaFold2-Multimer^80,81^ to predict potential Bero–AstC-R2 complex structures (see Methods). AlphaFold2-Multimer predicted an interaction between Bero and the extracellular region of AstC-R2 (Figures 6A and S12A). By contrast, the predicted Bero–AstC-R1 and Bero–5-HT1B complexes either had low confidence scores (AstC-R1, ranks 2–5; 5-HT1B, ranks 2 and 3) or positioned Bero at the cytoplasmic loops of these receptors (AstC-R1, rank 1; 5-HT1B, ranks 1, 4, and 5) (Figures S12B and S12C). The latter configurations are incompatible with the extracellular localization of GPI-anchored Bero.

Previous studies have shown that ligand properties can influence G protein coupling by GPCRs.^82^ For example, somatostatin receptor 2 (SSTR2) couples predominantly to Gαi/o upon binding somatostatin, whereas binding of the larger ligand octreotide has been associated with Gαq/11 coupling.^83^ Because Bero (104 amino acids) is considerably larger than the AstC peptide (15 amino acids), we hypothesized that Bero binding might favor an AstC-R2 conformation associated with Gαq/11 signaling. In the predicted Bero-bound AstC-R2 structure, several transmembrane helices were displaced outward relative to those in the predicted AstC-bound structure (Figure S13A), resembling structural differences reported for ligand-bound SSTR2 (FigureS13B).

Next, we investigated whether Bero and AstC-R2 physically associate in *Drosophila* Schneider 2 (S2) cells. Immunoprecipitation of FLAG-tagged Bero (Bero^FLAG^) co-precipitated HA- and GFP-tagged AstC-R2 (AstC-R2^HA&GFP^; see Methods), supporting a physical association between Bero and AstC-R2 when co-expressed in the same cells (Figures 6B and 6C). On immunoblotting, AstC-R2 was detected as a putative monomeric species with an apparent mass of approximately 75 kDa (triangle in Figures 6B, S14B, and S14C) and as an uncharacterized high-molecular-weight species (>250 kDa; arrow in Figures 6B, S14B, and S14C). Because the high-molecular-weight species was detected using two independent epitope tag-specific antibodies (anti-GFP in Figure 6B and anti-HA in Figure S14C), it is unlikely to represent an antibody-specific artifact. It may instead represent an SDS-resistant oligomer or another high-molecular-weight complex. Similar high-molecular-weight species have been reported for several GPCRs, including SSTR2 and the μ-opioid receptor, the mammalian homolog of AstC-R2.^84,85^ By contrast, Bero^FLAG^ did not detectably co-immunoprecipitate with 5-HT1B^GFP^ (Figures S14D, and S14E). Together, these biochemical results support the physical association between Bero and AstC-R2 when the proteins are co-expressed in the same cells.

Having established a physical association between Bero and AstC-R2, we next asked whether Bero indirectly facilitates AstC-R2 activation by the secreted AstC peptide or acts on AstC-R2 in *cis.* To address this question, we tested whether Bero and AstC-R2 could reconstitute a functional signaling module in type III neuroendocrine cells, which regulate adult wing expansion.^86,87^ Our reanalysis of scRNA-seq datasets indicated that type III neuroendocrine cells show no detectable expression of *bero*^34^ or *AstC-R2* (Figure S15A). Ectopic expression of *bero* alone allowed normal wing expansion in most flies, whereas expression of *AstC-R2* alone suppressed wing expansion in approximately half of the flies (Figures 4D and S15B). These results suggest that AstC-R2 may exhibit constitutive, ligand-independent signaling activity. Strikingly, ectopic co-expression of *bero* and *AstC-R2* in these cells completely suppressed wing expansion (Figures 4D and S15B), consistent with strong inhibition of the neuroendocrine output required for wing expansion. This effect was reproduced using the two independent GAL4-drivers for type III neuroendocrine cells, *burs-GAL4* and *CCAP-GAL4* (Figures S15B and S15C). Together, these data indicate that Bero and AstC-R2 can form a functional *cis*-acting signaling module capable of modifying neuronal output, even in a cellular context lacking detectable endogenous expression of *bero* and *AstC-R2*.

### Secreted AstC peptide counteracts Bero–AstC-R2 signaling under high osmolality

AstC peptides are the canonical endogenous inhibitory *trans*-ligands of AstC-R2 (Figure S16A),^88–90^ raising the question of how secreted AstC peptides regulate AstC-R2 when the receptor is engaged by membrane-tethered Bero in *cis*. We therefore hypothesized that osmosensitive AstC-producing neurons release AstC peptides under high-osmolality conditions, thereby suppressing Bero–AstC-R2 signaling in ABLK neurons. GFP-tagged AstC-R2 (AstC-R2^GFP^) expressed in LK neurons localized to neurites in close proximity to AstC-immunoreactive axons (Figures S16B and S16C), consistent with the possibility that AstC peptides act on ABLK neurites.

To investigate whether AstC peptides transmit a signal associated with hemolymph osmolality, we examined the osmosensitivity of ABLK neurons in an *AstC* loss-of-function mutant.^91,92^ In the *AstC* mutant, ABLK neuronal activity significantly increased in S-330 but not in S-280 (Figures 7A and B). Notably, loss of AstC increased ABLK activity in S-330 but did not fully restore it to the level observed under low-osmolality conditions (Figure 7B), suggesting that additional osmotic signals may converge on this circuit mechanism. These findings indicate AstC contributes substantially, but not exclusively, to the osmolality-dependent modulation of ABLK neuronal activity.

We next tested whether AstC peptides were sufficient to suppress ABLK neuronal activity by applying synthetic AstC peptides while monitoring intracellular Ca²⁺ levels. Previous studies have shown that synthetic AstC must undergo oxidation to form an intramolecular disulfide bond and become a functional ligand (Figure 7C).^88^ Application of oxidized AstC in S-280 caused a significant dose-dependent decrease in intracellular Ca²⁺ levels in ABLK neurons (Figures 7D, 7E and S17). By contrast, untreated AstC peptides did not reduce ABLK Ca²⁺ levels (Figures 7D, 7E and S17). Notably, AstC also reduced Ca²⁺ levels in the presence of Cd²⁺, which inhibits voltage-gated Ca²⁺ channels and blocks neurotransmitter release.^93^ These results suggest that AstC suppresses ABLK neuronal activity without requiring synaptic transmission.

**Figure 7.**
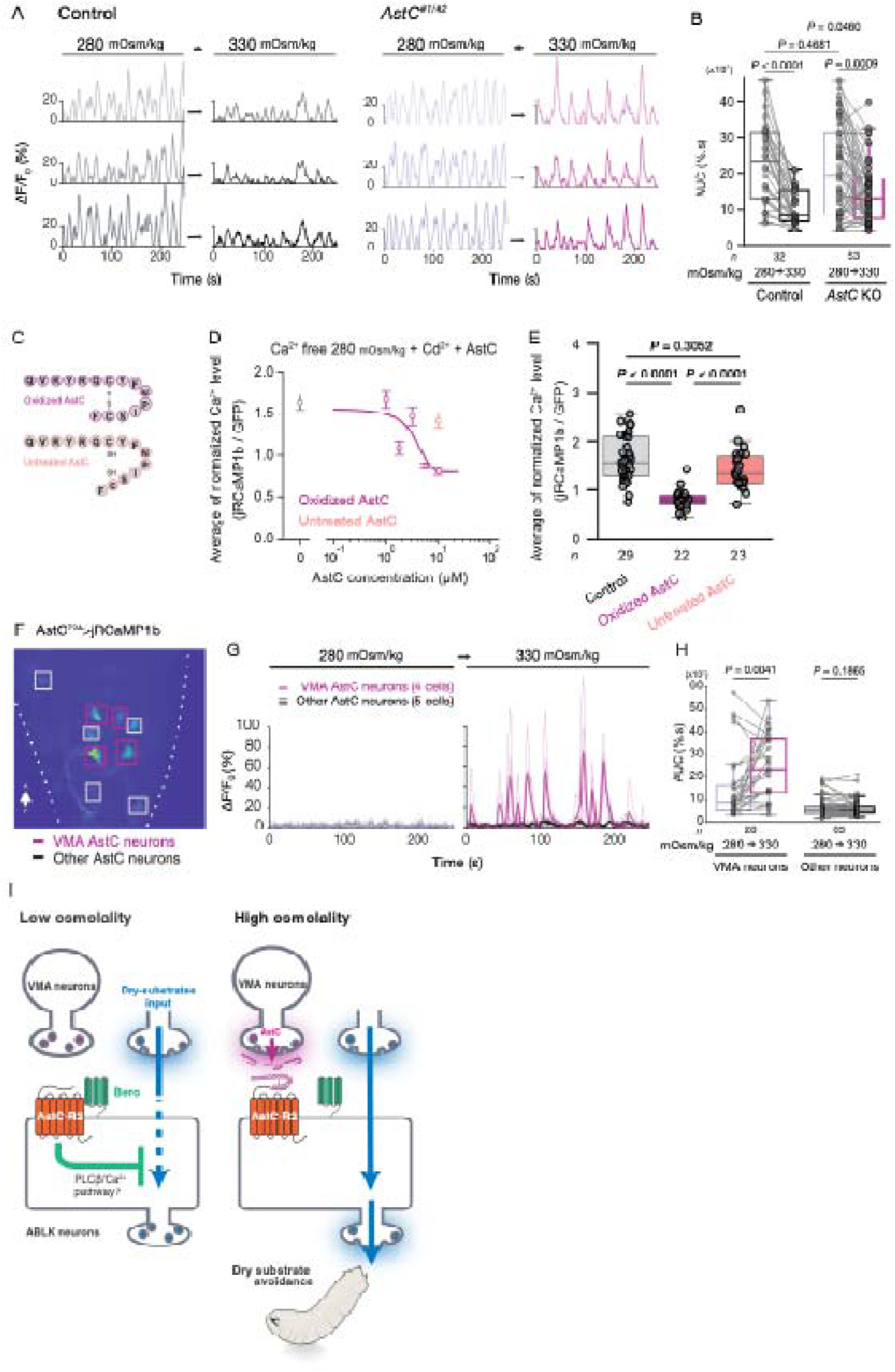
Secreted AstC peptide counteracts Bero–AstC-R2 signaling under high osmolality. (A) Representative traces of Ca²⁺ activity (Δ*F*_Persistent_/*F*_0_) in ABLK neurons under 280 and 330 mOsm/kg conditions in control and *AstC* knockout larvae. (B) Quantification of spontaneous Ca²⁺ activity, measured as area under the curve (AUC), in control and *AstC* knockout larvae. Control, *n* = 32 neurons; *AstC* knockout, *n* = 53 neurons. Wilcoxon rank-sum test for unpaired comparisons; Wilcoxon signed-rank test for paired comparisons. (C) Schematic representation of oxidized and untreated AstC peptides. (D and E) Effects of oxidized and untreated AstC peptides on intracellular Ca²⁺ levels in ABLK neurons. *n* = 20–29 neurons from 3–4 larvae. Data are shown as mean ± s.e.m. in (D), with quantification of normalized intracellular Ca²⁺ levels in (E). Kruskal–Wallis test, followed by Dunn’s test. (F to H) Ca²⁺ activity of AstC^+^ neurons in the ventral nerve cord (VNC) under different osmolality conditions. *n* = 91 neurons from 10 larvae, including 28 VMA-AstC neurons and 63 other AstC^+^ neurons. (F) Representative image showing Ca²⁺ levels in AstC^+^ neurons. (G) Representative traces of spontaneous Ca²⁺ activity (Δ*F*_Persistent_/*F*_0_) in VMA-AstC neurons and other AstC^+^ neurons under 280 and 330 mOsm/kg conditions. (H) Quantification of spontaneous Ca²⁺ activity, measured as AUC, in VMA-AstC neurons and other AstC^+^ neurons across osmolality conditions. Wilcoxon signed-rank test. (I) Model illustrating how Bero and AstC link internal osmotic state to avoidance behavior. See also Figure S16-S19.

It has been hypothesized that the AstC–AstC-R2 complex decreases intracellular Ca²⁺ levels via Gαi/o signaling, although direct experimental evidence is limited.^88,94^ However, the AstC-induced reduction in Ca²⁺ levels was not blocked by inhibition of Gαi/o signaling (Figure S18A). Thus, AstC does not appear to suppress Ca²⁺ activity by simply activating the canonical Gαi/o pathway; instead, it may interfere with coupling of the Bero–AstC-R2 complex to PLCβ signaling. To investigate this possibility, we performed co-immunoprecipitation experiments in the presence of AstC peptide. AstC treatment did not detectably alter the association between AstC-R2 and Bero (Figure S18B), suggesting that AstC does not suppress Bero–AstC-R2 signaling by disrupting complex formation. The mechanism by which AstC modulates signaling by the Bero–AstC-R2 complex therefore remains unclear.

Finally, we investigated whether AstC-producing neurons themselves are sensitive to osmolality (Figure S19A). A specific subpopulation of AstC^+^ neurons, previously termed VMA-AstC neurons,^89,95^ was selectively activated in S-330 (Figures 7F–H and S19B). These findings support a model in which high osmolality activates VMA-AstC neurons, leading to AstC-dependent suppression of ABLK neuronal activity.

## DISCUSSION

Animals maintain physiological homeostasis by continually adjusting their behavior according to internal states,^1^ and our findings reveal a non-canonical ligand–receptor mechanism by which the internal state of fluid osmolality reconfigures navigation behavior on dry substrates. At the circuit level, ABLK neurons act as a node at which the internal osmotic state, relayed through AstC^+^ neuronal activity, gates Class IV neuron-derived sensory transmission. The membrane-tethered Ly6/uPAR protein Bero interacts with the neuropeptide receptor AstC-R2 in *cis* to establish PLCβ-dependent Ca²⁺ activity in ABLK neurons. This high-Ca²⁺ activity state suppresses nociceptive transmission under low-osmolality conditions. Our data are consistent with a model in which AstC peptide released under high-osmolality conditions counteracts Bero–AstC-R2 signaling, thereby facilitating dry-substrate avoidance. Thus, membrane-tethered and secreted ligands can act in opposition on a shared GPCR signaling module, through which the organismal physiological state reshapes behavior.

A previous study showed that dry substrates pose a risk of failed pupation and death, but that larvae are able to avoid such substrates.^11^ However, it was unclear which behaviors enable this avoidance and how they are tuned by the internal state produced by desiccation stress. By experimentally separating prior desiccation experience from the immediate humidity conditions during navigation, we found that dry-substrate avoidance, facilitated by the internal dehydration state, guides larvae toward suitable pupation sites. During this behavior, raising the head and/or tail reduces the contact area between the dry surface and the larval body wall, facilitating detachment from the substrate and thereby causing the larva to fall (Figures 1 and S2). In their natural habitats, falling may carry larvae to relatively moist microhabitats that often lie vertically below, such as beneath fallen leaves (Figure S2C), and may therefore serve as an effective strategy for moving to a more favorable environment. Because holometabolous insects cannot relocate once they enter the immobile pupal stage, selecting a favorable pupation site while still mobile as larvae represents a primary—and perhaps the only—opportunity to mitigate the risk of pupal dehydration. Our data indicate that larvae use their internal osmotic state to preemptively select suitable pupation sites.

At the physiological level, our results link desiccation-induced increases in hemolymph osmolality to the enhanced dry-substrate avoidance through a defined neuropeptidergic pathway. Elevated extracellular osmolality activated VMA-AstC neurons, and AstC peptide suppressed ABLK neuronal activity. Because loss of AstC reduced, but did not abolish, the osmolality-dependent suppression of ABLK activity, AstC is likely one of several signals that convey osmotic state to this circuit node. This arrangement, in which a secreted neuropeptide modulates a constitutively active *cis*-ligand–receptor module, may allow internal-state signals to be integrated locally at a single interneuron class.

Our results indicate that an endogenous LU superfamily protein can act as a *cis*-ligand for a GPCR to modulate intracellular signaling and, consequently, animal behavior. This mechanism extends the conventional concept of ligand-dependent bias in GPCR signaling. In ABLK neurons, the Bero–AstC-R2 module elicits PLCβ-dependent Ca²⁺ activity, a signaling mode that has not previously been reported for AstC-R2 in any cell type. Given that LU superfamily proteins are widespread and many interact with transmembrane receptors,^23–26,96,97^ membrane-tethered ligands may be an unexplored source of diversity in endogenous GPCR signaling.

How PLCβ-dependent Ca²⁺ activity reduces nociceptive transmission remains to be determined.^33^ PLCβ cleaves phosphatidylinositol 4,5-bisphosphate (PIP₂), generating inositol 1,4,5-trisphosphate (IP₃) and diacylglycerol (DAG). Intracellular Ca²⁺ released from endoplasmic reticulum via the IP₃ receptor, together with DAG, could mediate protein kinase C (PKC)-dependent modulation of excitability, as previously described for SSTRs.^98,99^ This molecular-level mechanism may allow the same sensory cue to produce different behavioral outcomes according to internal needs without requiring circuit-wide alterations in sensory transmission. Together, these findings provide a molecular framework for understanding how internal osmotic state is translated into the opposing actions of membrane-tethered and secreted GPCR ligands, illustrating a broader principle by which animals flexibly adjust their behavior to maintain homeostasis under changing internal and external conditions.^1,15–19^

### Limitations of the study

This study has several limitations. First, the link between hemolymph osmolality and behavior rests partly on inference. We showed that desiccation raises hemolymph osmolality and enhances dry-substrate avoidance, and that osmolality modulates ABLK and VMA-AstC activity ex vivo. However, we did not manipulate hemolymph osmolality directly in behaving larvae. We also did not test whether manipulating AstC, AstC-R2, or VMA-AstC neurons alters HTR behavior or pupation-site selection. Second, the cells that directly sense changes in osmolality remain unidentified. The incomplete rescue of ABLK activity in the AstC mutant suggests that other osmotic signals also act on this circuit. Furthermore, our ex vivo saline solutions changed osmolality mainly through sucrose and trehalose and may not reproduce the full compositional changes in hemolymph that occur during desiccation. Third, the *cis*-ligand model is supported by structural predictions, co-immunoprecipitation from overexpressing S2 cells, and an ectopic-expression assay with an indirect behavioral readout. Direct structural or live-cell measurements of Bero– AstC-R2 engagement, and of its modulation by AstC peptide, are still needed to confirm this mechanism and the receptor conformational changes involved. Fourth, how AstC opposes Bero–AstC-R2 signaling remains unresolved. The G protein coupling of the complex was inferred from computational prediction and pharmacology (a PLCβ inhibitor and PTX), which do not clearly separate the contributions of Gαq/11, Gαi/o, and Gβγ. Finally, our behavioral experiments were conducted under controlled laboratory conditions, the field observations of HTR behavior were qualitative, and the agent-based model simplified substrate wetness and larval behavior. How strongly this mechanism shapes pupation-site choice in natural habitats therefore remains to be determined.

## RESOURCE AVAILABILITY

### Lead contact

Requests for further information and resources should be directed to and will be fulfilled by the lead contact, Tadao Usui.

### Materials availability

All unique *Drosophila* lines generated in this study are available from the lead contact.

### Data and code availability

• All data reported in this paper will be shared by the lead contact upon request.

• All original code has been deposited at Zenodo and is publicly available at https://doi.org/10.5281/zenodo.24122789.

• Any additional information required to reanalyze the data reported in this paper is available from the lead contact upon request.

## ACKNOWLEDGMENTS

We thank Mayumi Futamata, Kanae Oki, Hiroko Imai, Risa Muraki, and Yuriko Niitani for excellent technical assistance; members of the Uemura lab for experimental advice and discussions; Dr. Takaomi Sakai for the anti-LK primary antibody; Dr. Jan Adrianus Veenstra and Dr. Nilay Yapici for the anti-AstC primary antibody; Dr. Hiromu Tanimoto and Dr. Shu Kondo for the *AstC-R2-T2A-GAL4* strain; Dr. Craig Montell and Dr. Young-Joon Kim for the *AstC* mutant strains; Dr. Dai Watanabe and Dr. Satoshi Yawata for sharing the vapor pressure osmometer; and Mr. Takeshi Mita, Dr. Takahisa Ueno, Dr. Manabu Sekiguchi, and Dr. Airi Sato for valuable advice on the field survey. We also thank Dr. Hokto Kazama, Dr. Yoshitomo Kurogi, Dr. Eisuke Imura, Dr. Yuta Takada, Mr. Masa A. Shimazoe, Dr. Chihiro Fujiyabu and Dr. Tomoko Ohyama for valuable discussions and helpful comments. We thank the Bloomington Drosophila Stock Center, Kyoto Drosophila Stock Center, and Korea Drosophila Resource Center for providing fly stocks, As well as FlyBase and the Drosophila Genomics Resource Center.

## AUTHOR CONTRIBUTIONS

YT and TUsui designed the study. YT generated ABLK split-GAL4 lines and performed behavioral assays, except for the side-view recordings in Figures 1C and 1D, computer simulations, Ca²⁺ imaging, histological imaging experiments, and bioinformatic analyses. MK performed Ca²⁺ imaging experiments on SELK, A08n, and AstC neurons and measured hemolymph osmolality. TSO performed Ca²⁺ imaging experiments with L-glucose and biochemical experiments with assistance from RM and JS. HK recorded behavior from the side view with assistance from AN. KM investigated the conditions for exposure to desiccation stress with assistance from YT. EM performed immunostaining for AstC in fig. S16B. AT investigated saline conditions for Ca²⁺ imaging together with YT and KL. TUsui built the optogenetic systems and generated the constructs and transgenic flies. MK created the illustrations. YT and TUsui drafted the manuscript. TUsui supervised the study. All authors, including TUemura, participated in discussions and edited the manuscript.

## DECLARATION OF GENERATIVE AI AND AI-ASSISTED TECHNOLOGIES IN THE WRITING PROCESS

During the preparation of this work, the authors used ChatGPT (OpenAI) and Claude (Anthropic) for language editing and proofreading of the manuscript. After using this tool or service, the authors reviewed and edited the content as needed and take full responsibility for the content of the publication.

## SUPPLEMENTAL INFORMATION

Document S1. Supplemental methods, Figures S1–S19 and Videos S1-S7

## STAR METHODS

### KEY RESOURCES TABLE

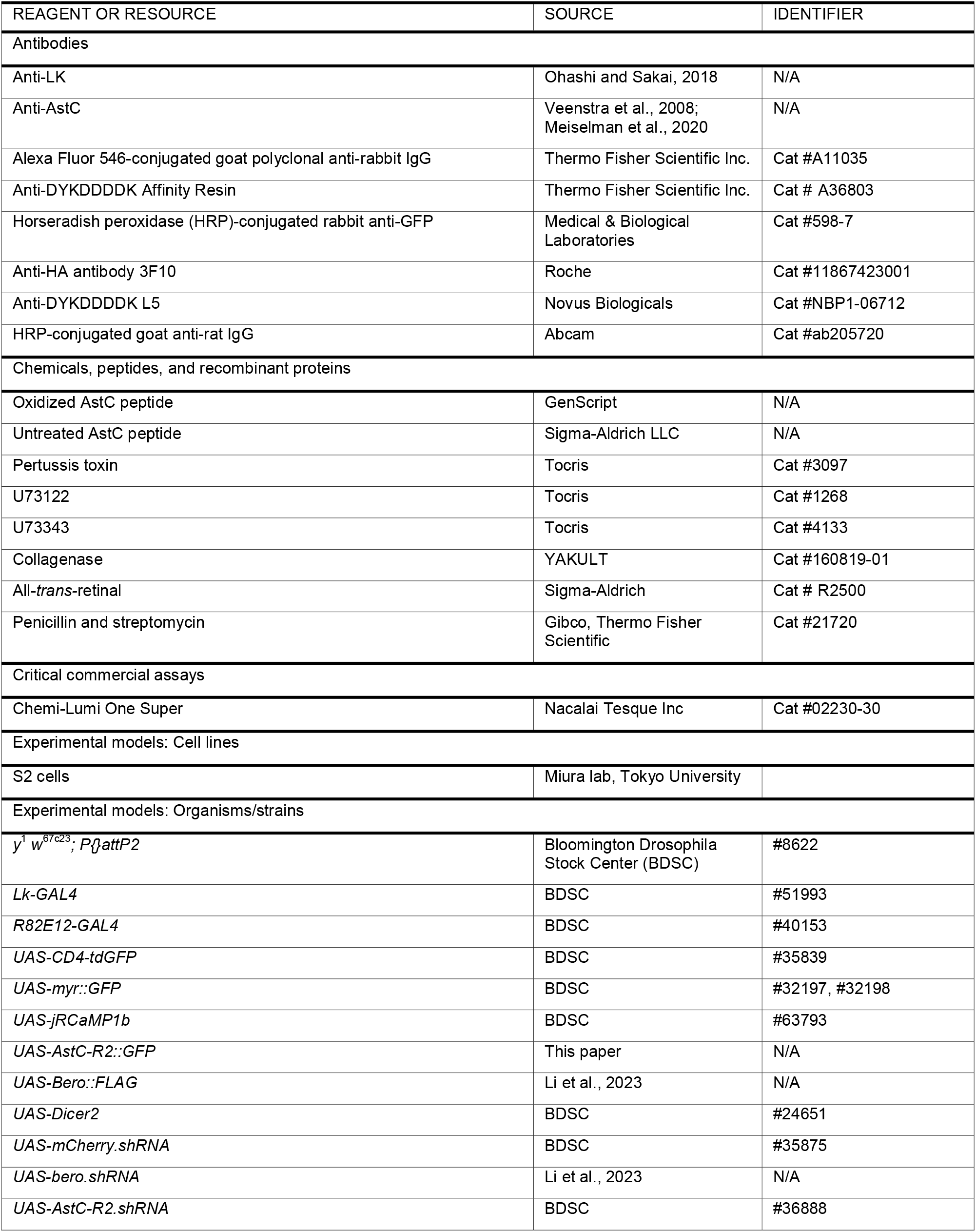
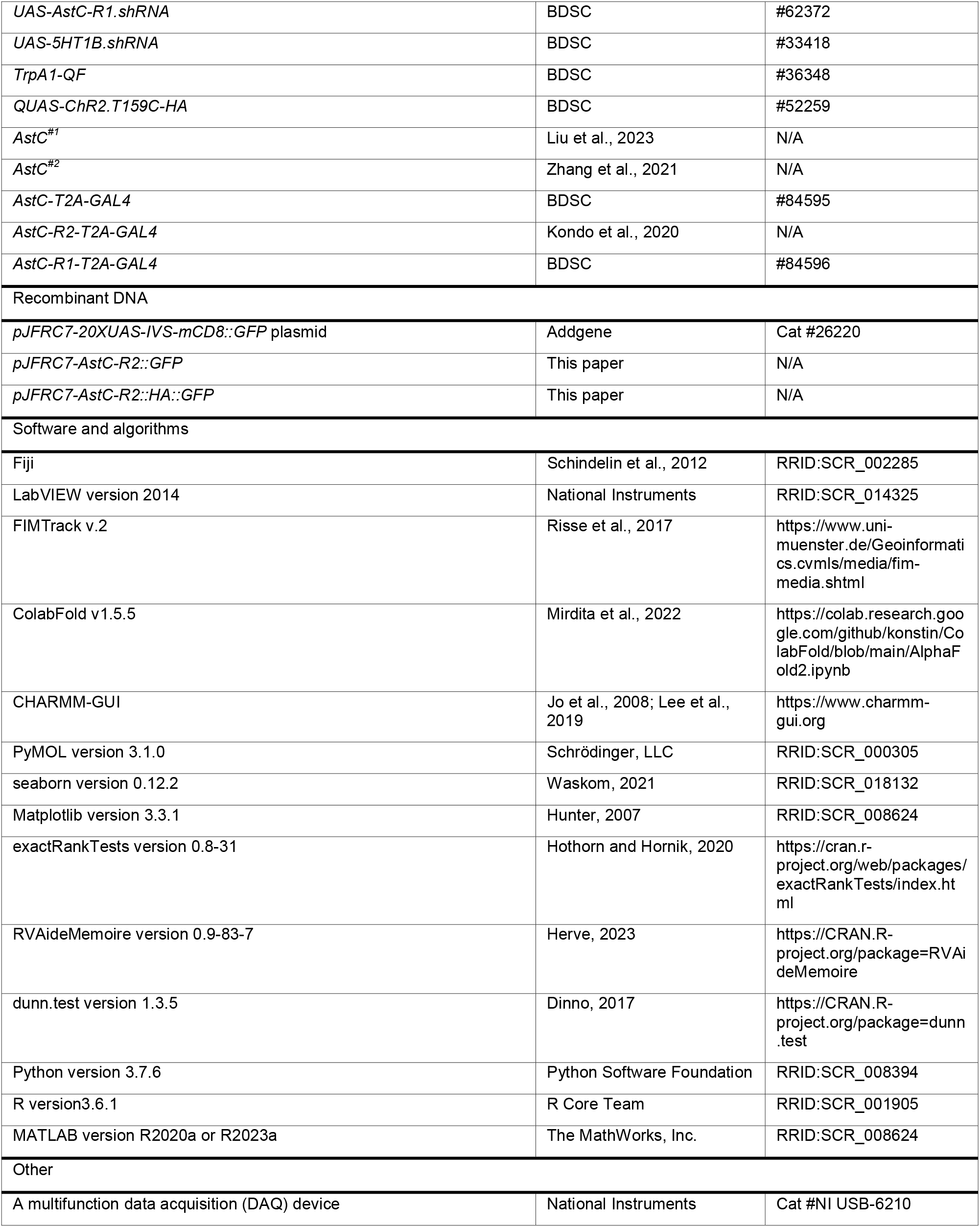

| REAGENT OR RESOURCE | SOURCE | IDENTIFIER |
|---|---|---|
| Antibodies |  |  |
| Anti-LK | Ohashi and Sakai, 2018 | N/A |
| Anti-AstC | Veenstra et al., 2008; Meiselman et al., 2020 | N/A |
| Alexa Fluor 546-conjugated goat polyclonal anti-rabbit IgG | Thermo Fisher Scientific Inc. | Cat #A11035 |
| Anti-DYKDDDDK Affinity Resin | Thermo Fisher Scientific Inc. | Cat # A36803 |
| Horseradish peroxidase (HRP)-conjugated rabbit anti-GFP | Medical & Biological Laboratories | Cat #598-7 |
| Anti-HA antibody 3F10 | Roche | Cat #11867423001 |
| Anti-DYKDDDDK L5 | Novus Biologicals | Cat #NBP1-06712 |
| HRP-conjugated goat anti-rat IgG | Abcam | Cat #ab205720 |
| Chemicals, peptides, and recombinant proteins |  |  |
| Oxidized AstC peptide | GenScript | N/A |
| Untreated AstC peptide | Sigma-Aldrich LLC | N/A |
| Pertussis toxin | Tocris | Cat #3097 |
| U73122 | Tocris | Cat #1268 |
| U73343 | Tocris | Cat #4133 |
| Collagenase | YAKULT | Cat #160819-01 |
| All-*trans*-retinal | Sigma-Aldrich | Cat # R2500 |
| Penicillin and streptomycin | Gibco, Thermo Fisher Scientific | Cat #21720 |
| Critical commercial assays |  |  |
| Chemi-Lumi One Super | Nacalai Tesque Inc | Cat #02230-30 |
| Experimental models: Cell lines |  |  |
| S2 cells | Miura lab, Tokyo University |  |
| Experimental models: Organisms/strains |  |  |
| *y<sup>1</sup> w<sup>67c23</sup>; P{}attP2* | Bloomington Drosophila Stock Center (BDSC) | #8622 |
| *Lk-GAL4* | BDSC | #51993 |
| *R82E12-GAL4* | BDSC | #40153 |
| *UAS-CD4-tdGFP* | BDSC | #35839 |
| *UAS-myr::GFP* | BDSC | #32197, #32198 |
| *UAS-jRCaMP1b* | BDSC | #63793 |
| *UAS-AstC-R2::GFP* | This paper | N/A |
| *UAS-Bero::FLAG* | Li et al., 2023 | N/A |
| *UAS-Dicer2* | BDSC | #24651 |
| *UAS-mCherry.shRNA* | BDSC | #35875 |
| *UAS-bero.shRNA* | Li et al., 2023 | N/A |
| *UAS-AstC-R2.shRNA* | BDSC | #36888 |
| *UAS-AstC-R1.shRNA* | BDSC | #62372 |
| *UAS-5HT1B.shRNA* | BDSC | #33418 |
| *TrpA1-QF* | BDSC | #36348 |
| *QUAS-ChR2.T159C-HA* | BDSC | #52259 |
| *AstC<sup>#1</sup>* | Liu et al., 2023 | N/A |
| *AstC<sup>#2</sup>* | Zhang et al., 2021 | N/A |
| *AstC-T2A-GAL4* | BDSC | #84595 |
| *AstC-R2-T2A-GAL4* | Kondo et al., 2020 | N/A |
| *AstC-R1-T2A-GAL4* | BDSC | #84596 |
| Recombinant DNA |  |  |
| *pJFRC7-20XUAS-IVS-mCD8::GFP* plasmid | Addgene | Cat #26220 |
| *pJFRC7-AstC-R2::GFP* | This paper | N/A |
| *pJFRC7-AstC-R2::HA::GFP* | This paper | N/A |
| Software and algorithms |  |  |
| Fiji | Schindelin et al., 2012 | RRID:SCR_002285 |
| LabVIEW version 2014 | National Instruments | RRID:SCR_014325 |
| FIMTrack v.2 | Risse et al., 2017 | https://www.uni-muenster.de/Geoinformatics.cvmls/media/fim-media.shtml |
| ColabFold v1.5.5 | Mirdita et al., 2022 | https://colab.research.google.com/github/konstin/ColabFold/blob/main/AlphaFold2.ipynb |
| CHARMM-GUI | Jo et al., 2008; Lee et al., 2019 | https://www.charmm-gui.org |
| PyMOL version 3.1.0 | Schrödinger, LLC | RRID:SCR_000305 |
| seaborn version 0.12.2 | Waskom, 2021 | RRID:SCR_018132 |
| Matplotlib version 3.3.1 | Hunter, 2007 | RRID:SCR_008624 |
| exactRankTests version 0.8-31 | Hothorn and Hornik, 2020 | https://cran.r-project.org/web/packages/exactRankTests/index.html |
| RVAideMemoire version 0.9-83-7 | Herve, 2023 | https://CRAN.R-project.org/package=RVAideMemoire |
| dunn.test version 1.3.5 | Dinno, 2017 | https://CRAN.R-project.org/package=dunn.test |
| Python version 3.7.6 | Python Software Foundation | RRID:SCR_008394 |
| R version3.6.1 | R Core Team | RRID:SCR_001905 |
| MATLAB version R2020a or R2023a | The MathWorks, Inc. | RRID:SCR_008624 |
| Other |  |  |
| A multifunction data acquisition (DAQ) device | National Instruments | Cat #NI USB-6210 |

### EXPERIMENTAL MODEL AND STUDY PARTICIPANT DETAILS

#### Drosophila melanogaster

*Drosophila melanogaster* strains were reared on standard fly food at 25°C and 75–80% relative humidity under a 12-h light/dark cycle, as previously described.^33^ Larvae used for optogenetic manipulation experiments were reared in the dark on fly food containing 0.5 mM all-*trans*-retinal (R2500; Sigma-Aldrich). Transgenic lines were maintained mainly in either white (w*) or yellow vermilion (y1, v1) mutant backgrounds, unless stated otherwise. The genotypes used in the experiments are shown in Table S1.

#### Drosophila S2 cells

S2 cells were maintained in Schneider’s Drosophila medium containing penicillin and streptomycin (21720; Gibco, Thermo Fisher Scientific) and supplemented with 10% fetal bovine serum (FBS).

### METHOD DETAILS

#### Generation of plasmids and transgenic strains

To generate the *UAS-AstC-R2::GFP* plasmid and transgenic strain, a genomic fragment containing the entire coding sequence of *AstC-R2* was amplified by polymerase chain reaction (PCR) from genomic DNA isolated from male *w*^11^^18^ flies. The *pJFRC7-20XUAS-IVS-mCD8::GFP* plasmid (#26220) was digested with *XhoI* and *BamHI* to remove the mCD8 sequence, and the PCR product was inserted into the digested vector using In-Fusion HD Cloning (Takara Bio Inc., Kusatsu, Japan). To generate the *UAS-AstC-R2::HA::GFP* plasmid, *pJFRC7-AstC-R2::GFP* was used as a template, and a 2×HA tag was inserted into the cytoplasmic-tail region of AstC-R2 using inverse fusion PCR cloning.

#### Immunohistochemistry and confocal imaging

To detect LK peptides in the larval CNS, wandering third-instar larvae were dissected in PBS and fixed overnight at 4°C in 4% formaldehyde in PBS. After fixation, the dissected CNS samples were washed five times with PBST (PBS containing 0.3% Triton X-100) and blocked for 30 min at room temperature in PBST containing 2% BSA filtered through a 0.22-µm filter. The samples were then incubated with rabbit polyclonal anti-LK antibody (1:1000)^100^ at 4°C for 3 days.

To detect AstC peptides, dissected CNS samples were fixed in 4% formaldehyde in PBS for 25 min at room temperature. After fixation, the samples were washed five times with PBST (PBS containing 0.2% Triton X-100) and blocked for 30 min at room temperature in PBST containing 5% normal goat serum (NGS). The samples were then incubated overnight at 23–25°C with rabbit polyclonal anti-AstC (1:500) antibody (1:500).^94,101^

After five washes with PBST, the samples were incubated with Alexa Fluor 546-conjugated goat polyclonal anti-rabbit IgG (H+L) antibody (1:500; A-11035, Thermo Fisher Scientific) for 2–3 h at room temperature. After additional washes, the samples were mounted using ProLong Glass Antifade Mountant (Thermo Fisher Scientific, Carlsbad, CA, USA). Images were acquired using a Nikon C1Si or Olympus FV3000 confocal microscope and processed using Fiji software (ImageJ, NIH, Bethesda, MD, USA).

The normalized values in fig. S16 were calculated by subtracting the minimum value from the signal within the ROI for each channel and dividing the difference by the maximum minus minimum value.

#### Pupation-site assay

Animals were reared in the dark at 25°C for 96–120 h. Wandering larvae were collected from the vials, briefly washed with deionized water, and transferred with a dry brush onto absorbent paper (KimWipes^®^, S200 62011 J-240, Nippon Paper Crecia Co., Ltd., Tokyo, Japan) to remove excess water (Figure 1A). To expose the larvae to either dry or humid conditions, they were transferred to empty plastic vials under the respective conditions. The vials were placed in sealed containers with silica gel or damp paper towels and left for 1 h. The larvae were then washed again with water, and excess water was removed. The larvae were carefully placed on the surface of fresh food. After more than 48 h at 25°C, the positions of the pupae were marked with oil-based ink and traced onto paper. The traced paper was scanned at a resolution of 6082 × 4300 pixels (ApeosPort-VIII C5573, FUJIFILM, Tokyo, Japan), and the height of each pupa from the food surface was measured using ImageJ software. Pupation heights were comparable between males and females (Figure S1B); therefore, subsequent experiments were performed using only female larvae.

#### Behavioral analysis on dry substrate: dry-surface assay

As in the pupation-site assay, female wandering larvae were exposed to either dry or humid conditions, washed with water, and blotted on KimWipes to remove excess water from the body wall. Using a wet brush, the larvae were placed on a dry 100 × 100 mm acrylic plate and left for more than 1 min to allow them to initiate self-righting and normal locomotion before recording began. When larvae with completely dry body surfaces were placed on a dry substrate, they often failed to complete self-righting^102,103^ and instead exhibited HTR-like movements. Therefore, we carefully avoided completely drying the larval body surface before placing the larvae on the dry plastic plate. A custom software program written in LabVIEW (version 2014; National Instruments, Austin, TX, USA) and a multifunction data acquisition (DAQ) device (NI USB-6210; National Instruments) were used to record larval behavior. Behavioral parameters were acquired using FIMTrack v.2.^104^ Optogenetic illumination systems were used as previously described.^33^

To detect HTR behavior automatically and in a high-throughput manner, we developed a machine-learning-based classification system (fig. S3A). A total of 36,000 video frames showing 19 larvae moving on dry surfaces and two larvae moving on wet surfaces were used for decision-tree training. The learning algorithms were compared using a smaller number of test samples, and a decision tree model was selected for further analysis. To determine the presence or absence of HTR behavior in a specific frame, 21 features extracted using FIMTrack v.2, together with the corresponding values from the preceding and following 15 frames, were used as inputs to the decision-tree model, yielding 651 parameters per frame. A total of 6,300 video frames showing three larvae on dry surfaces and one larva on wet surfaces were used for hyperparameter tuning by a grid search. The change in spine length two frames earlier was ultimately selected as the feature that most accurately classified the presence or absence of HTR behavior (fig. S3B). The classification accuracy on the test data was 96% (fig. S3C).

#### Recording of larval behavior in the field

In June 2024 and June–July 2025, we observed the behavior of wild larvae on the Faculty of Medicine Campus, Kyoto University, Kyoto, Japan (35.023°N, 135.777°E and 35.025°N, 135.778°E, respectively). On sunny days, we inspected areas around fallen ripe plum fruits and recorded the behavior of wild wandering *Drosophila* sp. larvae using an Olympus Tough TG-6 digital camera.

#### Agent-based model simulation

The induction probability parameters for behavioral states on dry surfaces were obtained from the dry surface assay. The corresponding parameters for the wet surface were obtained from our previous study. ^33^ We also incorporated a previously reported effect^11^ whereby wandering larvae wet a dry surface with water carried on their body surface when moving from a wet surface to a dry surface. The simulation was terminated at the 1,200th step, at which point most simulated larvae reached pupation heights comparable to those observed experimentally. To estimate the probability of falling (*P*_falling_) during HTR behavior, we recorded the behavior of larvae in vertically positioned vials for approximately 30–60 s and measured the time until each larva fell. Because most larvae in our experiments stopped moving approximately 6 h after the start of the experiment, we assumed that the 1,200th step corresponded to 6 h and estimated *P*_falling_ to range from 1.0 × 10^−1.9^ to 1.0 × 10^−1.5^.

#### Measurement of hemolymph osmolality

Body fluids were collected from control and desiccation-experienced female larvae in a temperature-controlled room at 4°C, as previously described.^64^ The osmolality of 10 µL of the collected body fluid was measured using a VAPRO 5520 osmometer (Wescor Inc., Logan, UT, USA).

#### Calcium imaging

Female wandering larvae were dissected in a Ca²⁺-free external saline solution (120 mM NaCl, 3 mM KCl, 4 mM MgCl₂, 10 mM NaHCO₃, 5 mM TES, 10 mM HEPES, and 10 mM glucose) at 25 ± 2°C. The osmolality was adjusted to 280 mOsm/kg with water, and the final pH was adjusted to 7.25 using NaOH. Poly D-lysine-coated glass-bottom dishes (D11130H; Matsunami Glass Ind., Ltd., Kishiwada, Japan) were filled with 200 µL of oxygenated imaging saline solution containing 1.5 mM CaCl₂ (Table S2), and the isolated CNS was attached to the dish. Cell bodies were imaged using the red fluorescent Ca²⁺ indicator jRCaMP1b^105^ at one frame per second with an EM-CCD image sensor (iXon X3 or iXon EM+DU-888; Andor Technology Ltd., Belfast, UK) at 25 ± 2°C, as previously described.^33^ After 250 s of imaging, the original saline solution was replaced three times with saline solutions of different osmolalities (Table S2).

For optogenetic activation of nociceptive neurons, the body wall attached with the CNS was pinned with insect pins to a handmade glass-bottom magnet plate. The blue light-gated cation channel ChR2.T159C^106^ was expressed in Class IV neurons. At 250 s after the start of imaging, 470-nm blue light was applied at 0.37 mW/mm² for 2.5 s using a TTL-controlled light-emitting diode (LED) driver system (M470L5 and LEDD1B; ThorLabs, Newton, NJ, USA) for optogenetic activation. The data were analyzed with a custom program written in MATLAB (version R2020a or R2023a; The MathWorks, Inc., Natick, MA, USA), as previously described^33^.

For pharmacological experiments involving 10 µM AstC peptides [oxidized AstC peptide was synthesized by GenScript as previously described,^90^ and untreated AstC peptide was synthesized by Sigma-Aldrich] or drugs [1 ng/µl pertussis toxin (3097, Tocris), 1µM U73122 (1268, Tocris), or 1 µM U73343 (4133, Tocris)], the CNS was treated with collagenase (160819-01; YAKULT) to increase tissue permeability to the peptide or drug, as previously described.^37,107^ Briefly, 0.5 mg/mL collagenase in Ca²⁺-free saline solution was applied for 30 s. The collagenase was washed out twice with a Ca²⁺-free saline solution containing the peptides or drugs. After 10 min, imaging was performed for 100 s. To inhibit neural network activity mediated by synaptic transmission, AstC peptides were applied together with Cd²⁺, which inhibits voltage-gated Ca²⁺ channels and thereby exocytosis.^93^ The concentrations of AstC peptides and drugs were determined on the basis of previous studies.^90,108^ The jRCaMP1b and tdGFP signals were acquired alternately within each 1-s interval. The average normalized Ca²⁺ level was calculated by dividing the jRCaMP1b fluorescence intensity by the GFP fluorescence intensity for each frame and averaging the resulting ratios over 100 frames.

#### Identification of genes expressed in ABLK neurons

Previously published scRNA-seq data from the third-instar larval CNS (GSE135810)^69,70^ were analyzed. First, preliminary clustering was performed on all cells expressing *Lk*, which identified two major neuronal clusters: one contained a larger number of cells, whereas the other co-expressed *ITP* and *sNPF*. We annotated the former cluster as “ABLK neurons” and the latter as “ALK neurons”. Highly variable genes were then extracted from both clusters and generate heatmaps of the single-cell gene expression profiles.

#### Protein structure prediction

The structures of Bero, AstC-R2, and the Bero–AstC-R2 complex were predicted using ColabFold v1.5.5.^80,81^ Cell membrane modeling was performed using CHARMM-GUI^109,110^ (https://www.charmm-gui.org). The three-dimensional structures were visualized using PyMOL (Version 3.1.0.).

#### Immunoprecipitation and biochemical analysis

S2 cells were maintained in Schneider’s *Drosophila* medium containing penicillin and streptomycin (21720; Gibco, Thermo Fisher Scientific) and supplemented with 10% fetal bovine serum (FBS). For co-immunoprecipitation, cells were transiently co-transfected with pDA (actin5C-GAL4) and the indicated pJFRC7-based plasmids.^111^ After 4 days, cell pellets (3 × 10 cells) were lysed on ice for 30 min in UltraRIPA buffer (F015B, BioDynamics Laboratory Inc.) containing complete protease inhibitor (Roche) and sonicated as previously described.^112^ For input sample, the supernatant from the cell lysates were recovered by centrifugation at 10,000 × *g* for 30 min at 4°C. For co-immunoprecipitation (Co-IP), Pierce Anti-DYKDDDDK Affinity Resin (A36803, Thermo Scientific) was used to pull down Bero^FLAG^. In the peptide competition assay, whole-cell extracts were incubated with 100 µM AstC before the addition of Anti-DYKDDDDK Affinity Resin. For Western blotting, horseradish peroxidase (HRP)-conjugated rabbit anti-GFP antibody (1:1,000; 598-7, Medical & Biological Laboratories), rat anti-HA antibody (3F10; 1:1,000; 11867423001, Roche), and rat anti-DYKDDDDK epitope tag antibody (L5; 1:2,000; NBP1-06712, Novus Biologicals) were used. HRP-conjugated goat anti-rat IgG antibody (1:20,000; ab205720, Abcam) was used as the secondary antibody. Signals were developed using a chemiluminescence assay kit (Chemi-Lumi One Super; 02230-30, Nacalai Tesque Inc., Japan), and detected using an ImageQuant LAS 4000 system (Cytiva, USA).

### QUANTIFICATION AND STATISTICAL ANALYSIS

Western blot signals were quantified using the Gel Analysis function in ImageJ. Regions of interest (ROIs) were defined from the top of the gel to approximately the 70-kDa position (Figure S14A). For each ROI, the baseline was determined by connecting the signal intensities at the upper and lower boundaries of the ROI (Figure S14B). Signal intensity was quantified as the area under the curve after baseline subtraction. Relative affinity was calculated by dividing the quantified co-IP signal by the corresponding input signal, as previously described.^113^

The sample sizes, statistical tests, and p-values for each dataset are indicated in the figures or in the figure legends. Statistical analyses were performed in R (version 3.6.1). Comparisons among more than two groups were performed using the Kruskal–Wallis test, followed by Dunn’s test with Benjamini– Hochberg correction for multiple comparisons, using the dunn.test package (version 1.3.5). Adjusted P values are reported for post hoc comparisons.

Box plots and violin plots were generated using seaborn (version 0.12.2). The center line indicates the median, the box limits indicate the 25th and 75th percentiles (interquartile range, IQR), and the whiskers extend to the most extreme data points within 1.5 × IQR from the box limits. Individual data points are overlaid on each box plot. In violin plots, the white dot indicates the median, the thick inner bar indicates the interquartile range, and the outline represents the kernel density estimate of the data distribution.

## SUPPLEMENTARY FIGURES

**Figure S1.**
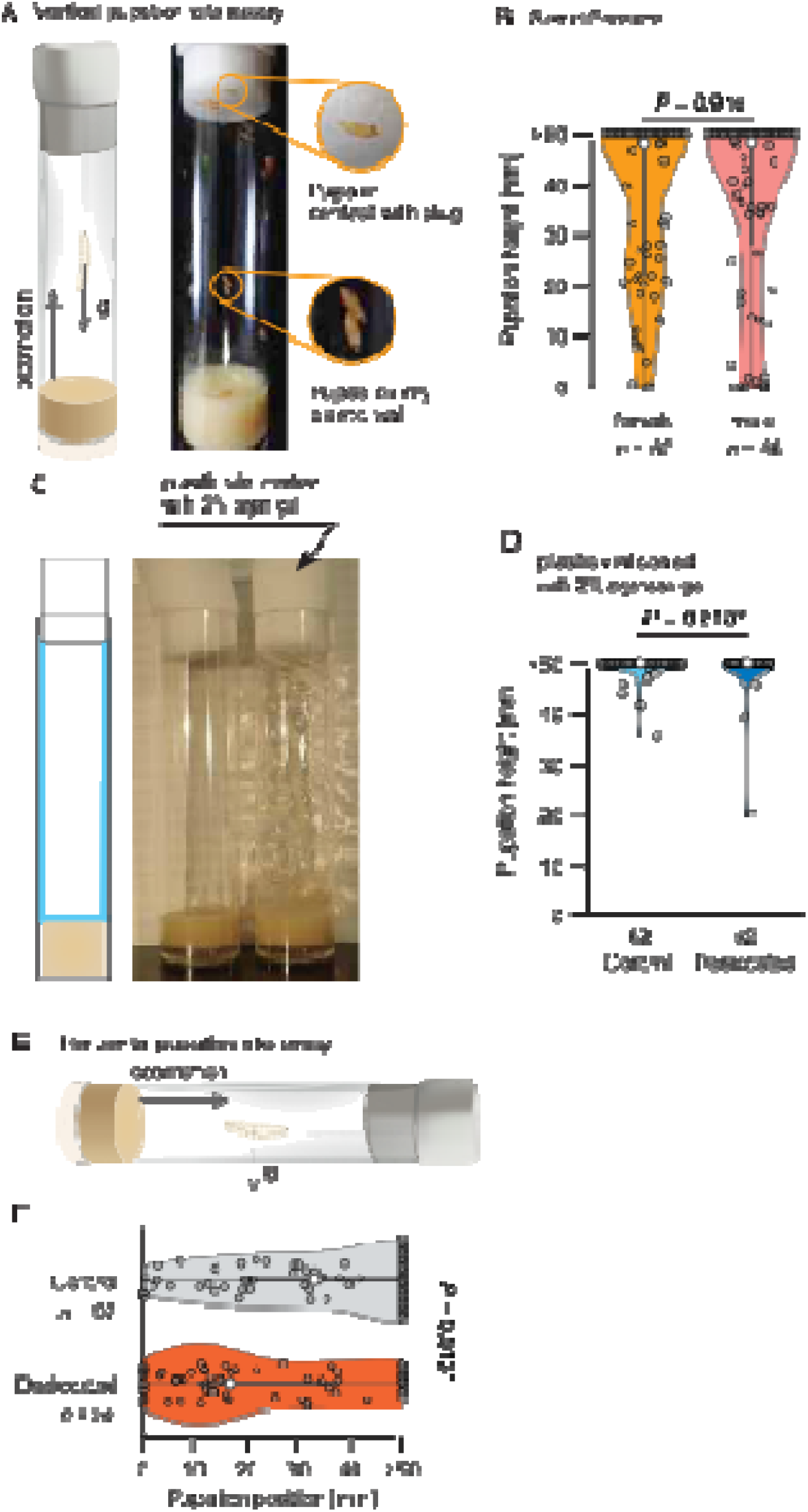
Dehydration modulates pupation-site navigation without impairing the climbing ability of *Drosophila melanogaster*. (A) Schematic of the vertical pupation-site assay. The downward arrow labeled “g” indicates the direction of gravity. Larvae that crawled upward often pupated at the junction between the plug and the vial wall. (B) Quantification of pupation height in male and female wandering larvae. Wilcoxon rank-sum test. (C) Schematic of the vertical pupation-site assay with the inner surface of the vial coated with 2% agar gel. (D) Quantification of pupation height in control and desiccation-experienced larvae in vertical vials with the inner wall coated with 2% agar. (E) Schematic of the horizontal pupation-site assay. The downward arrow labeled “g” indicates the direction of gravity. (F) Quantification of pupation position in the horizontal pupation-site assay in control and desiccation-experienced larvae. Wilcoxon rank-sum test.

**Figure S2.**
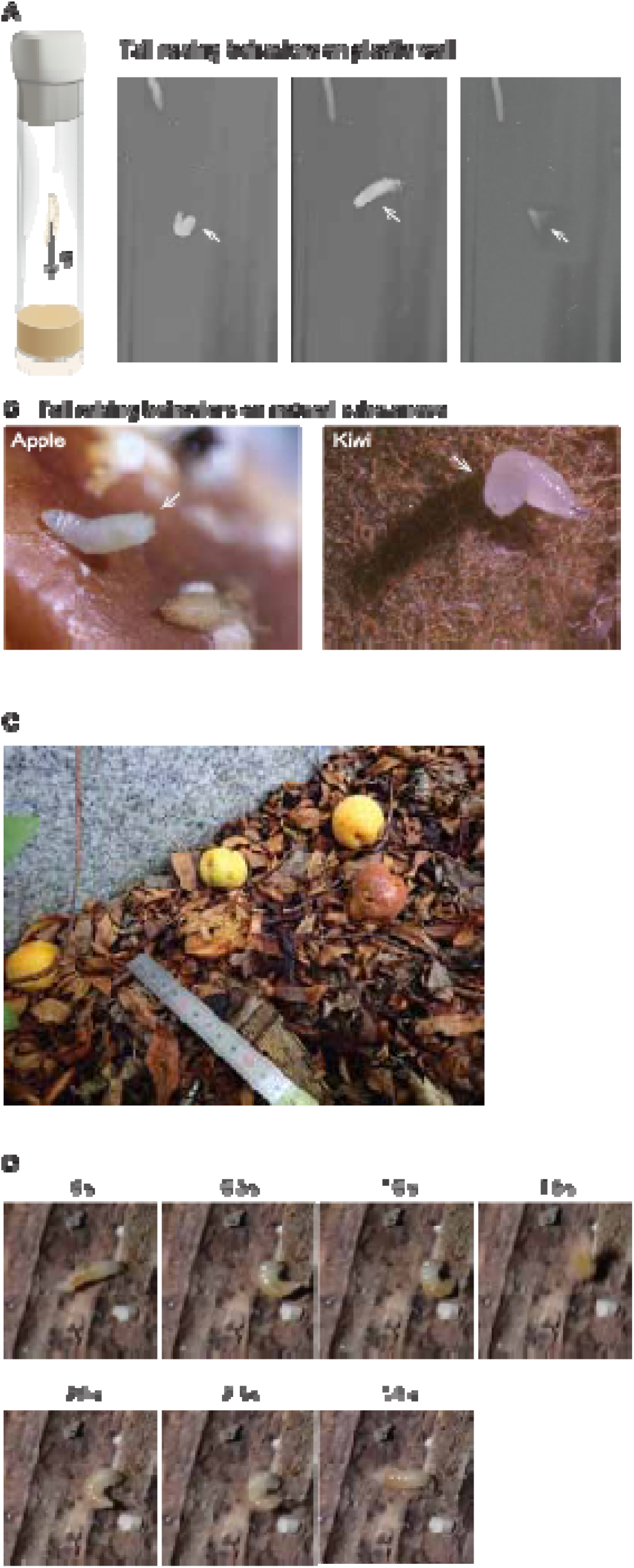
Head- and/or tail-raising behavior on various dry substrates. (A) Representative frames of head- and/or tail-raising (HTR) behavior on a plastic wall. Arrows indicate tail-raising larvae. (B) Representative images of HTR behavior on dry fruit surfaces. Arrows indicate tail-raising larvae. (C) Field site on the Kyoto University campus showing ripe plums scattered on the ground. (D) Representative frames of HTR behavior on the dry surface of wood chips.

**Figure S3.**
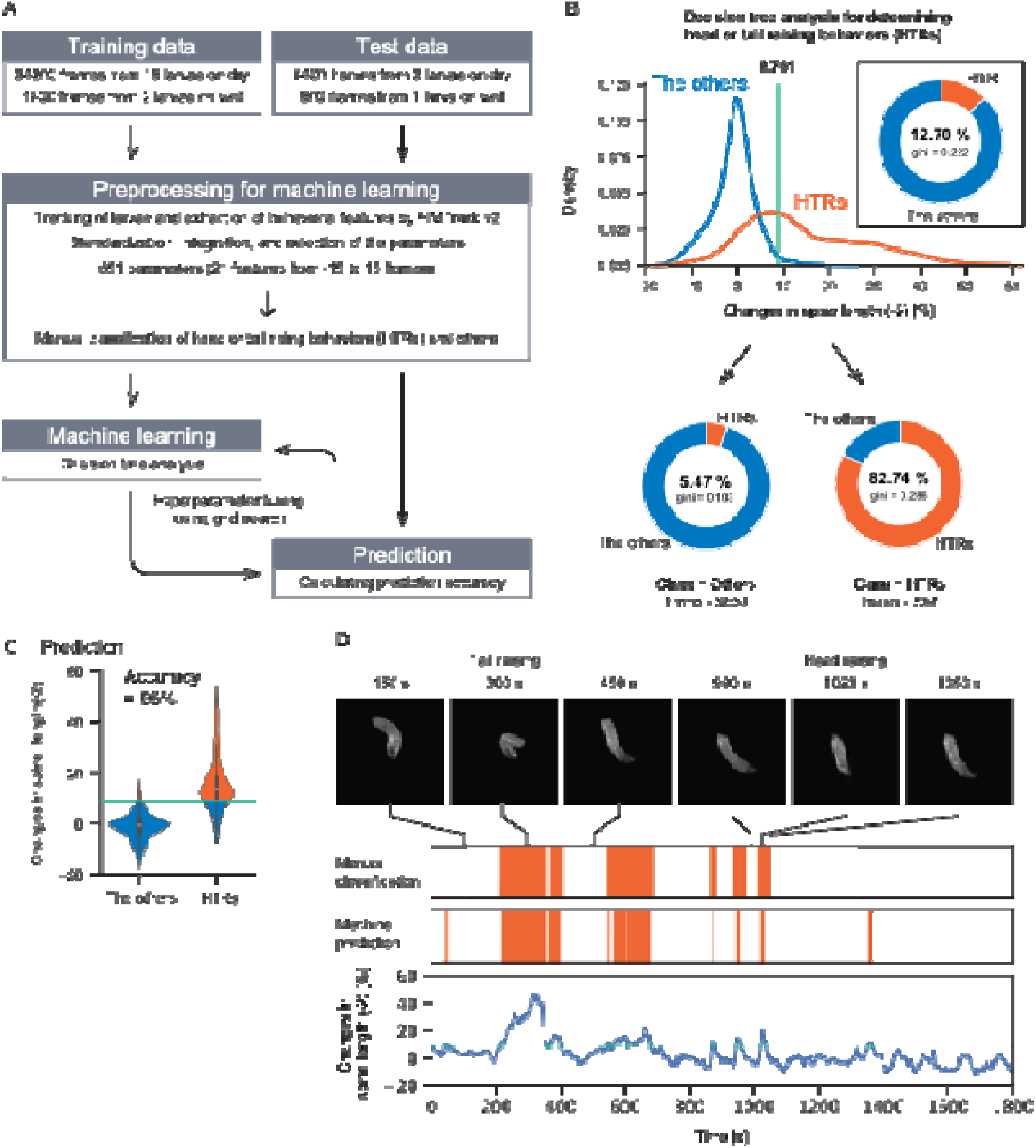
Development of a machine learning-based behavioral classifier for detecting HTR behavior. (A) Flowchart showing the training and evaluation of the decision tree classifier. Parameters were selected to maximize classification accuracy on the test data. (B) The feature that classified the training data with the highest accuracy was the percentage change in spine length two frames earlier [spine length change (–2)]. Using a threshold of 8.761 for spine length change (–2), frames containing HTR behavior, which accounted for 12.70% of the training data, were enriched to 82.74% after classification. Among frames classified as non-HTR, 5.47% contained HTR behavior (Gini coefficient = 0.103). (C) Classification accuracy for HTR behavior evaluated using the test dataset was 96%. (D) Top, representative frames of larvae on dry substrates. Middle, orange indicates frames in which HTR behavior was detected by manual annotation or the classifier. Bottom, time-series changes in spine length change (−2).

**Figure S4.**
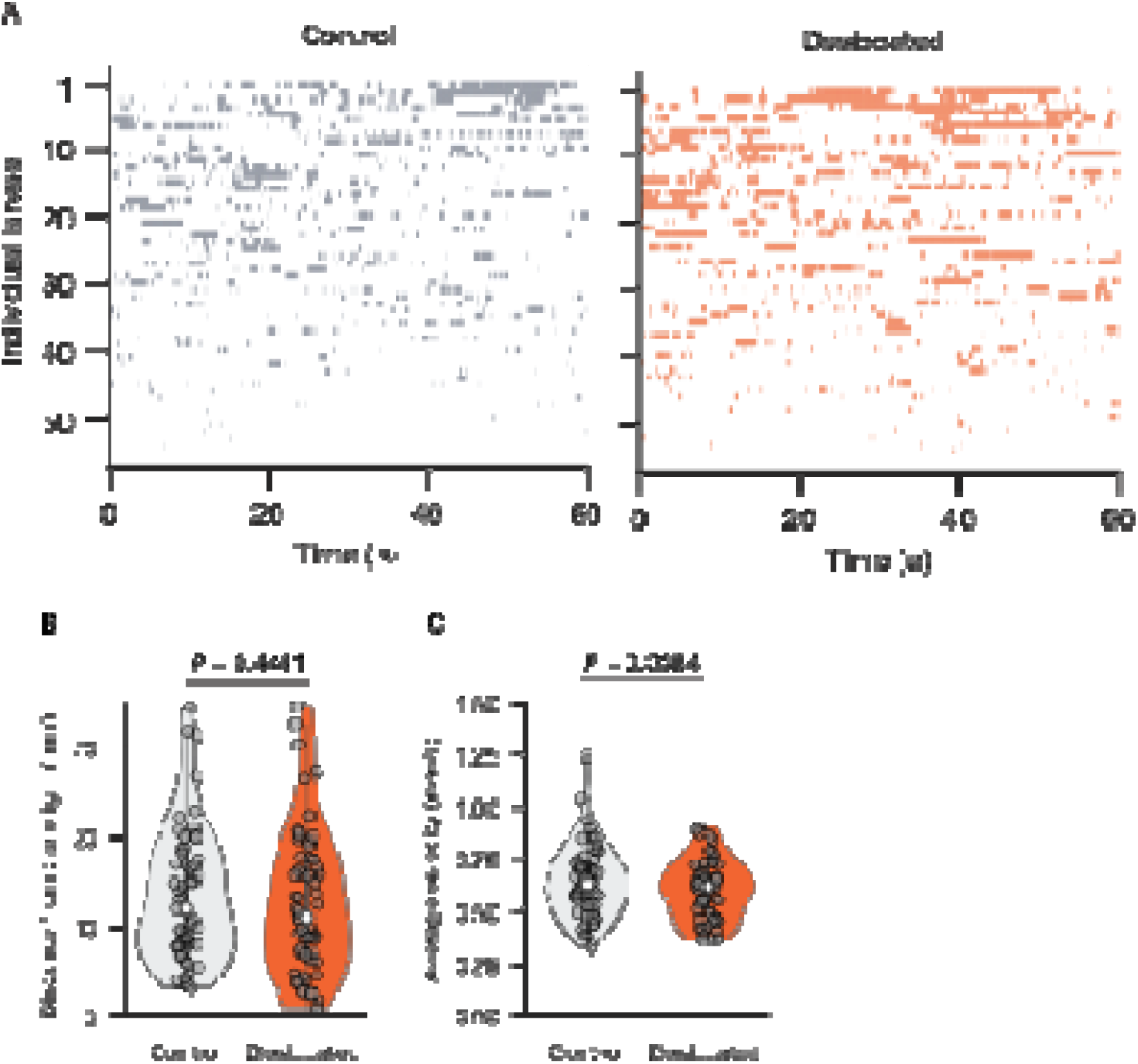
Dehydration promotes dry-substrate avoidance behavior. (A) Raster plots showing head- and/or tail-raising (HTR) behavior on dry substrate surfaces (control, *n* = 55; desiccation-experienced, *n* = 57). (B and C) Quantification of larval behavior on dry substrate surfaces during 1-min recordings: straight-line distance from the origin (B) and average velocity (C).

**Figure S5.**
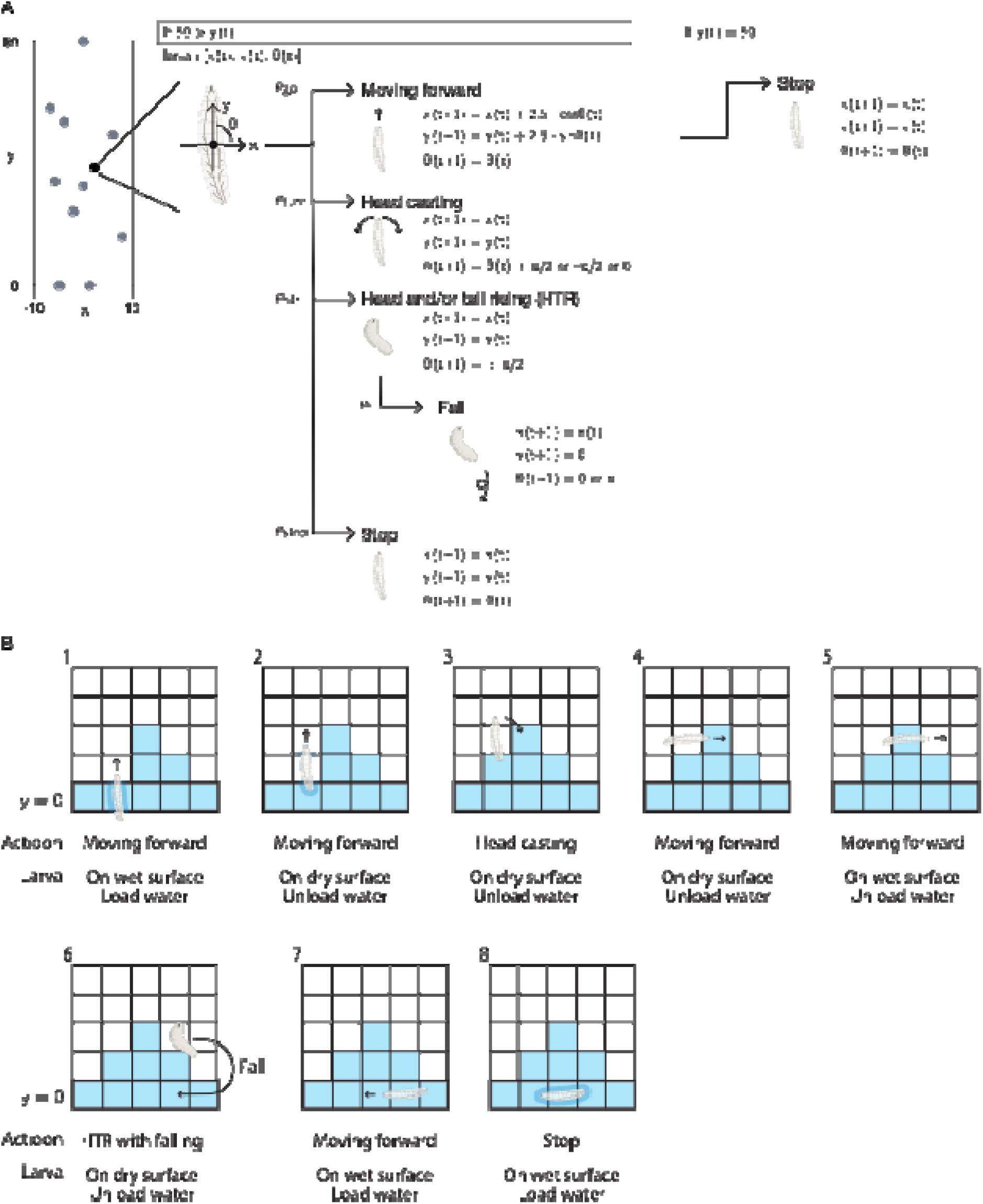
Schematic of an agent-based model simulating pupation-site selection. (A) The *i*th larval agent is represented by its position [*x* (*t*), *y* (*t*)] and body orientation, θ (*t*). At each time step, the larva selects a behavioral state according to the surface condition (wet or dry) and predefined transition probabilities. Possible actions include moving forward, head casting, head- and/or tail-raising (HTR), falling, and stopping. The selected action determines the position and orientation of the larva at the next time step. When the larva reaches the uppermost edge of the arena, it stops. (B) Example time series of larval movement in a grid-based environment. Blue grids indicate wet areas, and white grids indicate dry areas. The larva loads water while on a wet surface (indicated by blue shading around the larva in steps 1, 2, and 8) and unloads water after moving onto a dry surface. The white larval body represents the agent, and arrows indicate the direction of movement or changes in body orientation.

**Figure S6.**
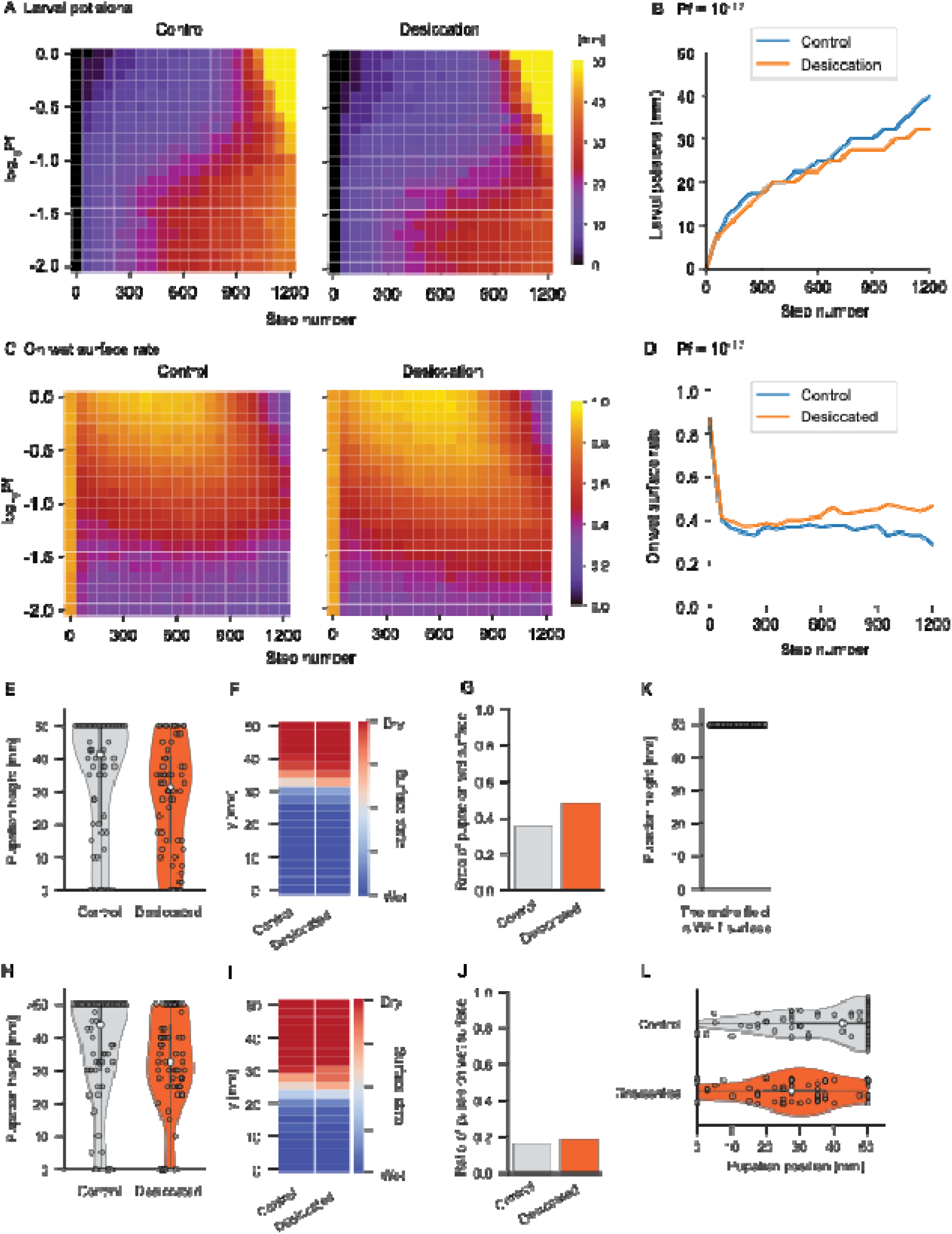
Results of the agent-based model simulation. (A) Representative heat maps showing the time-dependent distribution of simulated larval positions in vertical arenas under control and desiccation-experienced conditions. Color intensity indicates the relative density of larval positions during the simulation. White lines indicate the experimentally estimated range of falling probabilities during head- and/or tail-raising (HTR) behavior. (B) Time course of simulated larval positions extracted from the heat maps in (A) at *P*_falling_ = 1.0 × 10^−1.7^. (C) Heat maps showing time-dependent changes in simulated surface wetness in vertical arenas under control and desiccation-experienced conditions across the same range of *P*_falling_ values as in (A). Color intensity indicates the estimated level of surface wetness generated by larval movement over time. White lines indicate the experimentally estimated range of falling probabilities during HTR behavior. (D) Time course of simulated surface wetness extracted from the heat maps in (C) at *P*_falling_ = 1.0 × 10⁻¹·. (E) Quantification of simulated pupation height in control and desiccation-experienced larvae. The simulation was performed at *P*_falling_ = 1.0 × 10⁻¹· and recapitulated the experimental results shown in Fig. 1B. (F) Estimated spatial distribution of wet and dry surface regions at the end of the simulation at *P*_falling_ = 1.0 × 10⁻¹· . Blue and red indicate wet and dry surfaces, respectively. (G) Estimated proportion of larvae pupating on wet surface regions at *P*_falling_ = 1.0 × 10⁻¹· . (H) Quantification of simulated pupation height in control and desiccation-experienced larvae under a hypothetical condition in which larvae did not exhibit HTR behavior. (I) Estimated spatial distribution of wet and dry surface regions at the end of the simulation under a hypothetical condition in which larvae did not exhibit HTR behavior. Blue and red colors indicate wet and dry surfaces, respectively. (J) Estimated proportion of larvae pupating on wet surface regions under a hypothetical condition in which larvae did not exhibit HTR behavior. (K) Quantification of simulated pupation height in vertical vials with the inner wall surfaces coated with 2% agar gel. (L) Quantification of simulated pupation position in control and desiccation-experienced larvae in horizontal vials.

**Figure S7.**
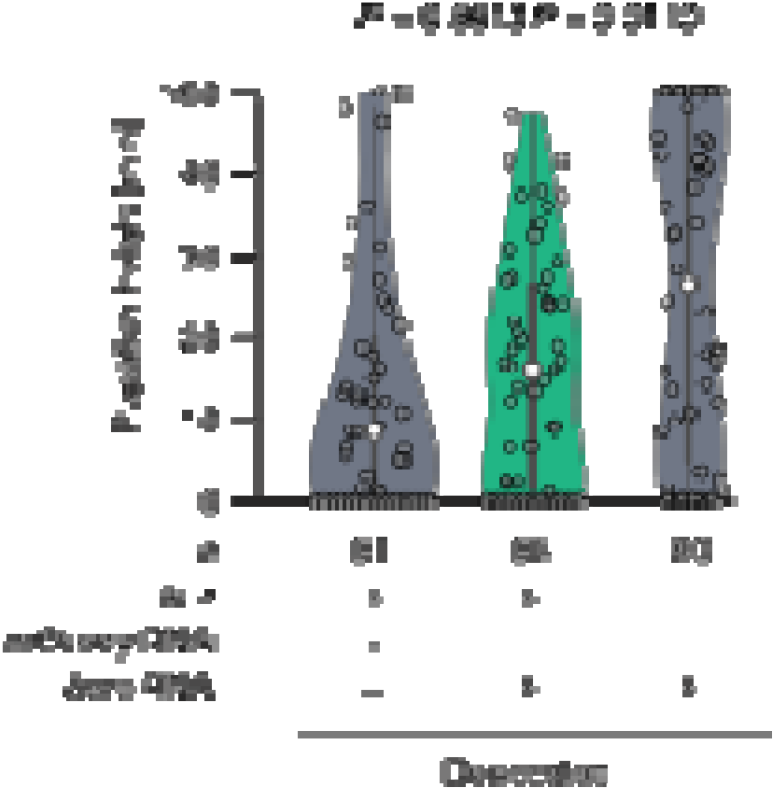
Pupation sites of desiccation-experienced larvae with *bero* RNAi. Pupation height of desiccation-experienced control larvae (LK > *mCherry* RNAi or + > *bero* RNAi) and desiccation-experienced larvae with *bero* knocked down in LK neurons (LK > *bero* RNAi). Kruskal–Wallis test followed by Dunn’s test.

**Figure S8.**
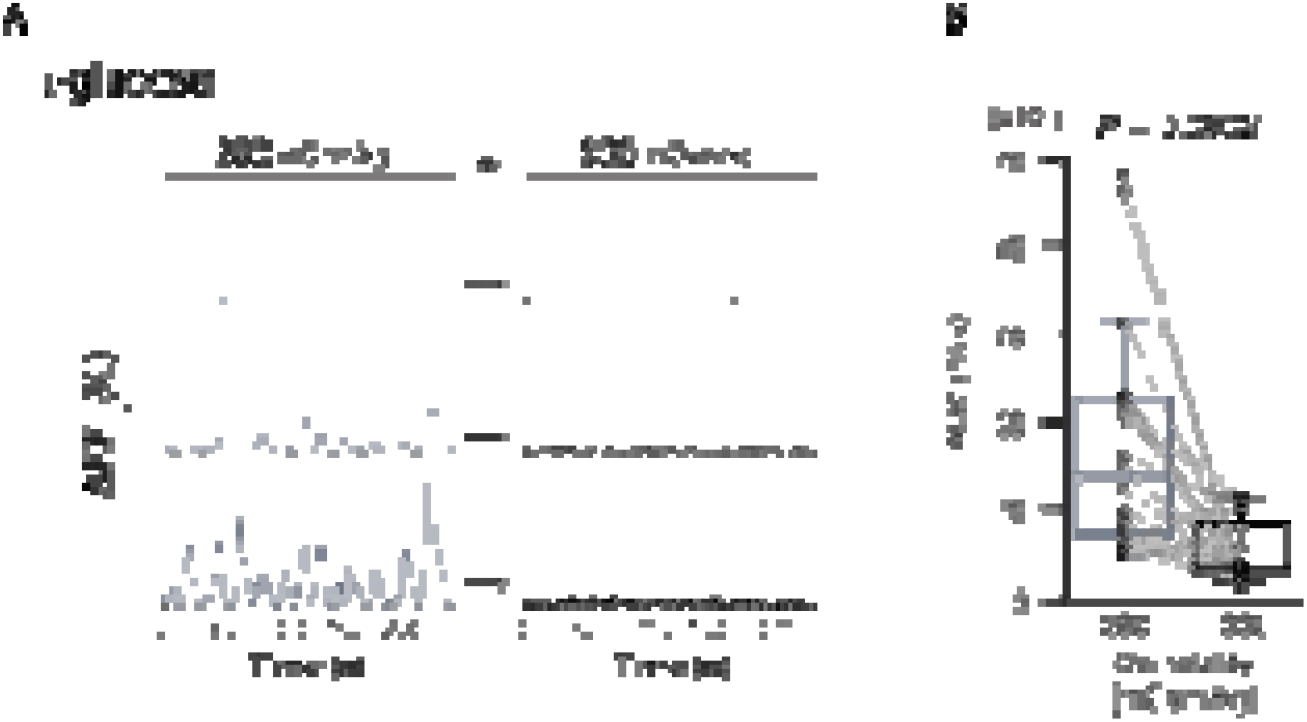
Osmosensitive neural activity of ABLK neurons. (A) Representative traces of spontaneous fluctuating Ca²⁺ activity (Δ *F*_Persistent_/*F*_0_) in ABLK neurons during sequential exposure to saline solutions of different osmolalities adjusted using the nonmetabolizable osmolyte L-glucose. (B) Quantification of spontaneous Ca²⁺ activity, measured as AUC, under different osmolality conditions adjusted using L-glucose (*n* = 20 neurons from four larvae).

**Figure S9.**
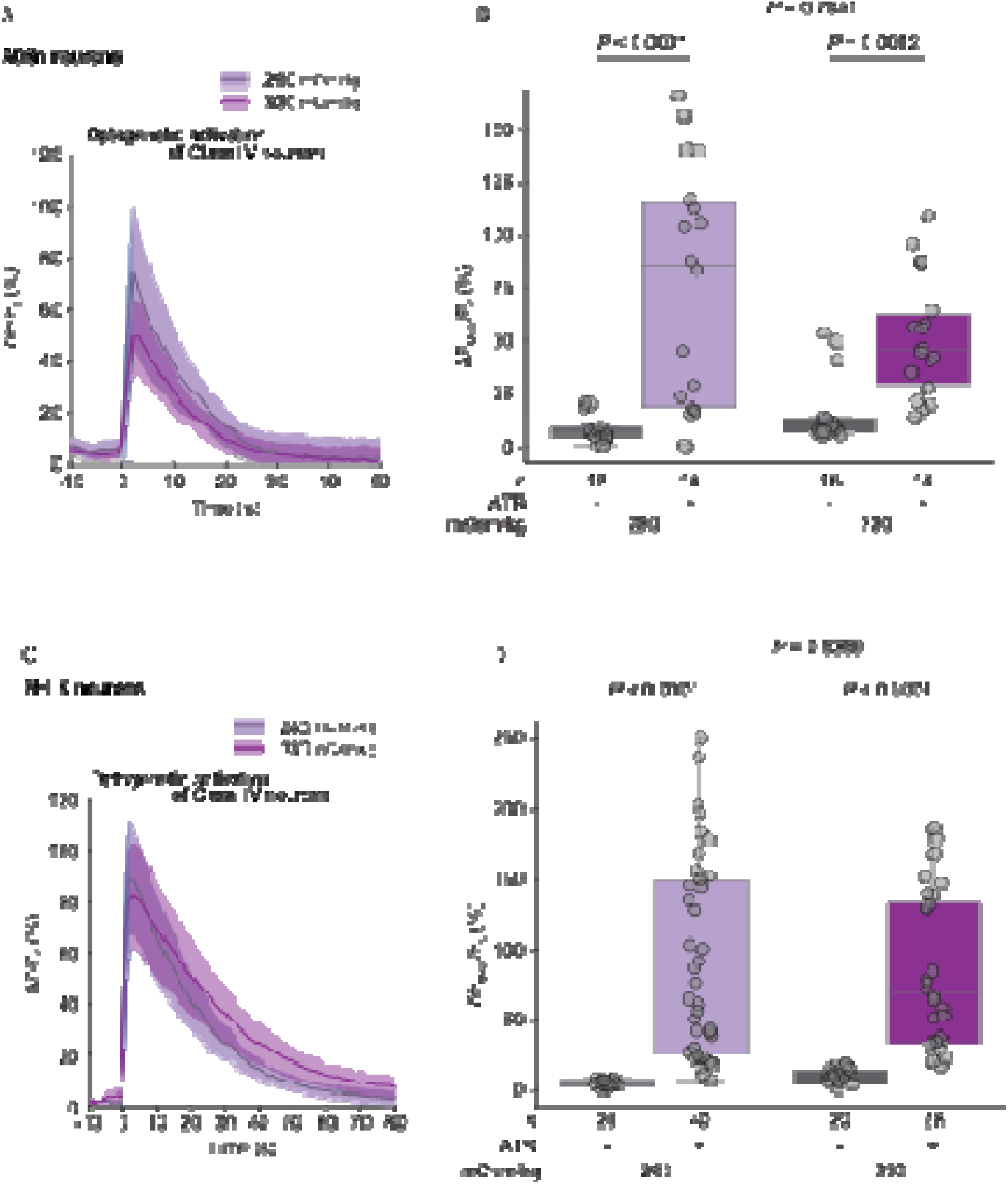
A08n and SELK neurons do not exhibit osmolality-dependent modulation of evoked sensory responses. (A) Representative evoked Ca²⁺ responses of A08n neurons following optogenetic activation of Class IV neurons. (B) Quantification of the maximum evoked Ca²⁺ response in A08n neurons. Wilcoxon rank-sum test. (C) Representative evoked Ca²⁺ responses of SELK neurons following optogenetic activation of Class IV neurons. (D) Quantification of the maximum evoked Ca²⁺ response in SELK neurons. Wilcoxon rank-sum test.

**Figure S10.**
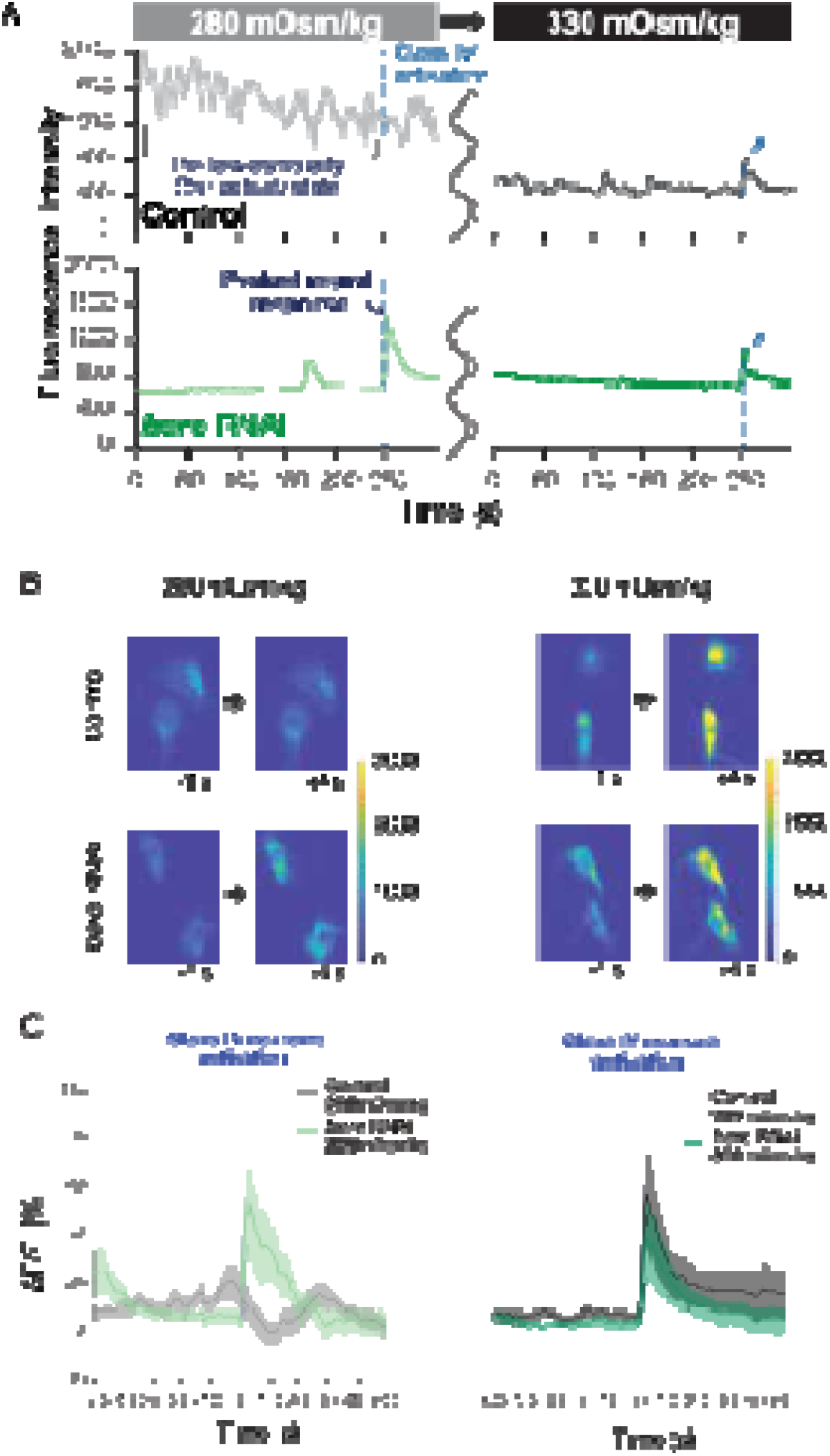
Bero modulates spontaneous activity and evoked nociceptive responses. (A) Representative traces of spontaneous fluctuating Ca²⁺ activity (jRCaMP1b fluorescence intensity) in ABLK neurons during sequential exposure to saline solutions with osmolalities of 280 and 330 mOsm/kg. (B) Representative confocal images of Ca²⁺ levels in AstC⁺ neurons in control and *bero* knockdown larvae under different osmolality conditions. (C) Representative evoked Ca²⁺ responses following optogenetic activation of Class IV neurons. Panels are related to Figure 4.

**Figure S11.**
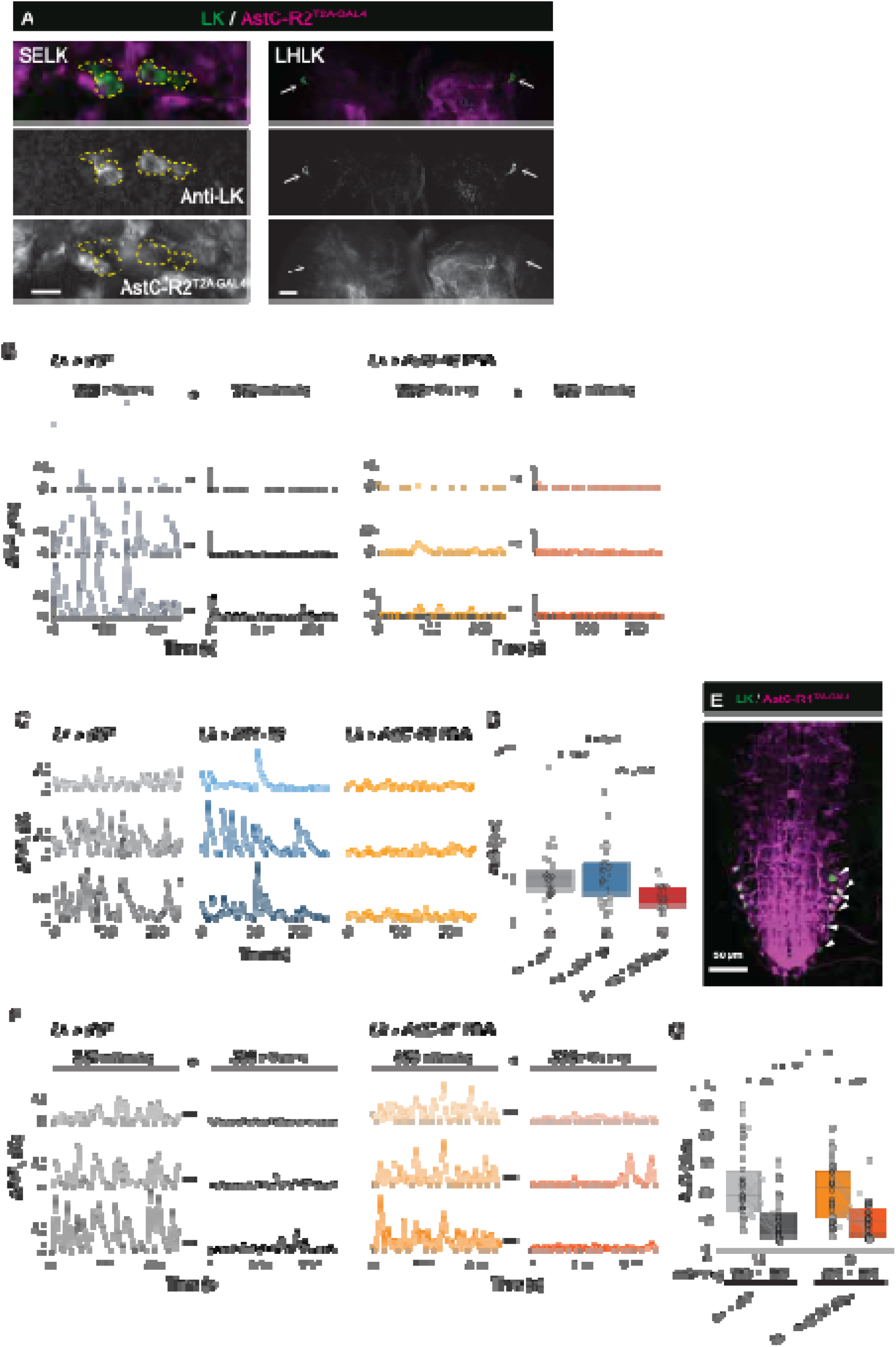
Identification of receptors involved in the osmolality-dependent activity of ABLK neurons. (A) Confocal images showing *AstC-R2-T2A-GAL4*-labeled cells (magenta) and SELK or LHLK neurons (green). Scale bars, 10 μm. (B) Representative traces of spontaneous fluctuating Ca²⁺ activity (Δ *F*_Persistent_/*F*_0_) in control and *AstC-R2* knockdown larvae under different osmolality conditions. (C) Representative traces of spontaneous fluctuating Ca²⁺ activity (Δ *F*_Persistent_/*F*_0_) in control, *5-HT1B* knockdown, and *AstC-R2* knockdown larvae at 280 mOsm/kg. (D) Quantification of spontaneous Ca²⁺ activity, measured as AUC. Kruskal–Wallis test, followed by Dunn’s test. (E) Confocal image showing *AstC-R1-T2A-GAL4*-labeled cells (magenta) and ABLK neurons (green). Scale bar, 50 μm. (F) Representative traces of spontaneous fluctuating Ca²⁺ activity (Δ *F*_Persistent_/*F*_0_) in control and *AstC-R1* knockdown larvae under different osmolality conditions. (G) Quantification of spontaneous Ca²⁺ activity, measured as AUC, in control and *AstC-R1* knockdown ABLK neurons at 280 and 330 mOsm/kg. Wilcoxon rank-sum test for unpaired comparisons and Wilcoxon signed-rank test for paired comparisons.

**Figure S12.**
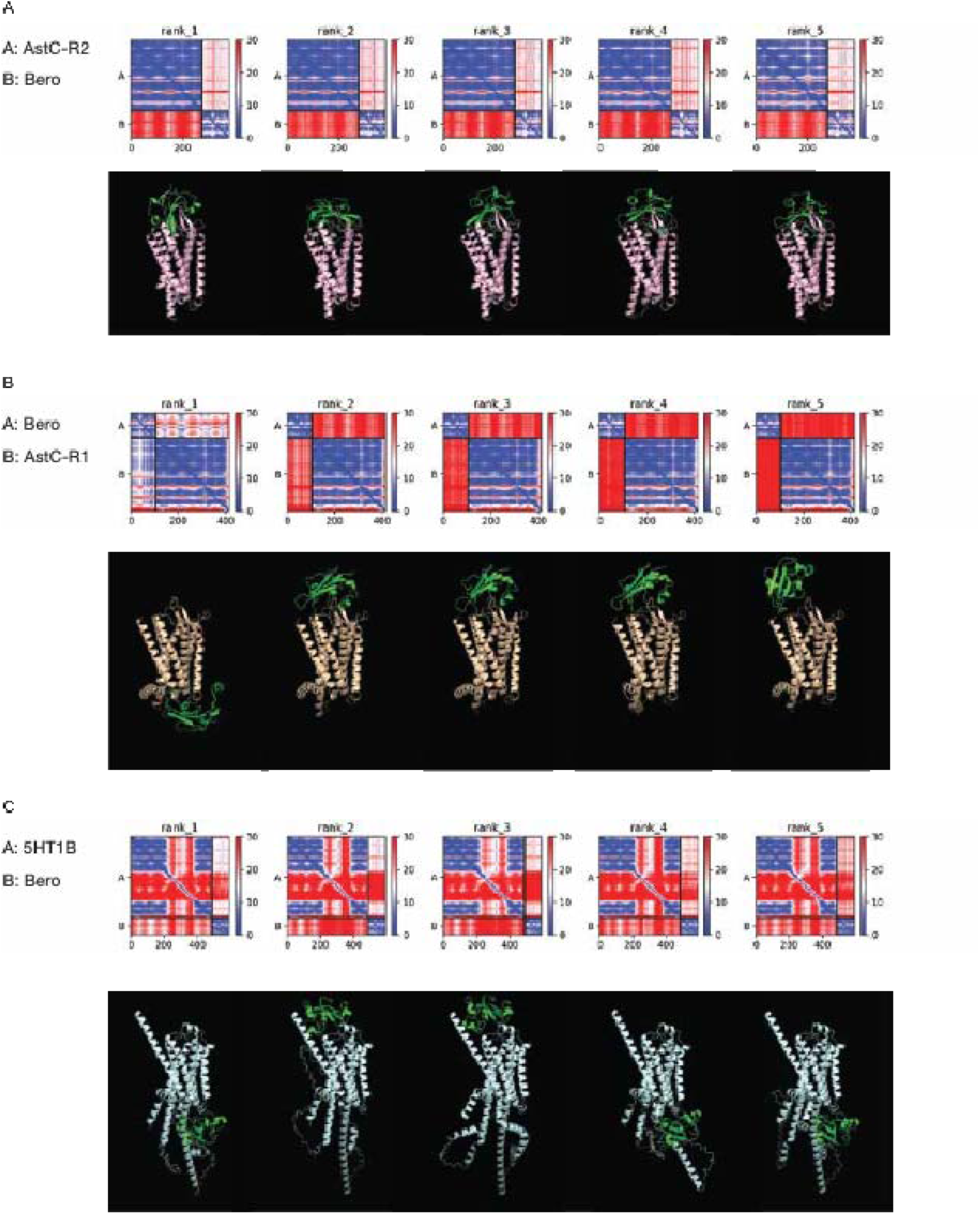
Prediction of Bero–GPCR complex structures. (A to C) Bero–GPCR complex structures predicted using AlphaFold2-Multimer. Predicted aligned error plots (top) and corresponding predicted complex structures (bottom) are shown for Bero with AstC-R2 (A), AstC-R1 (B), and 5-HT1B (C).

**Figure S13.**
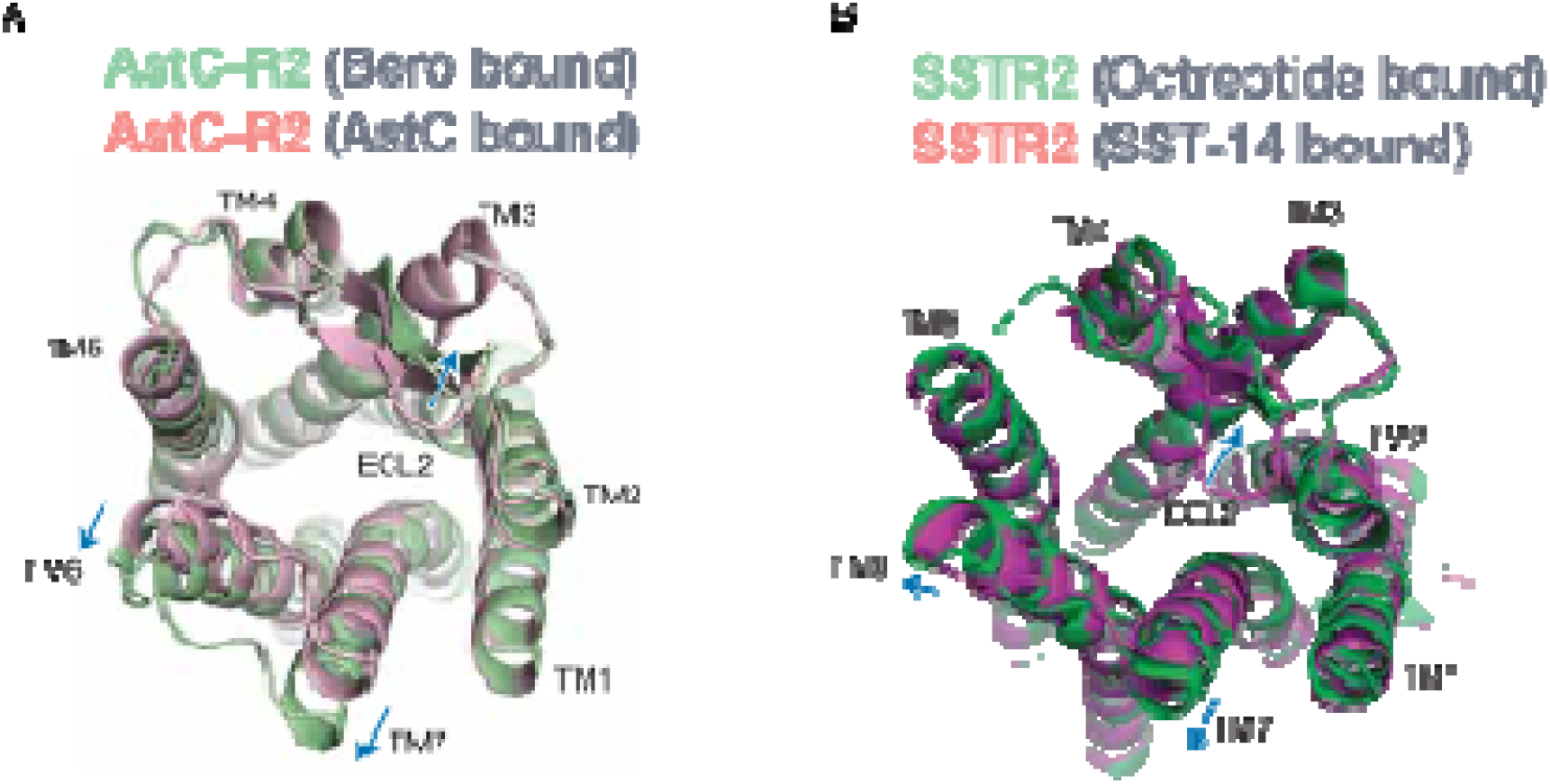
Structural comparison of predicted AstC-R2 complexes and SSTR2 structures. (A) Structural overlay of predicted Bero-bound and AstC-bound AstC-R2 structures generated using AlphaFold2. (B) Structural overlay of octreotide-bound and SST-14-bound SSTR2 structures (PDB IDs: 7Y26 and 7Y27).

**Figure S14.**
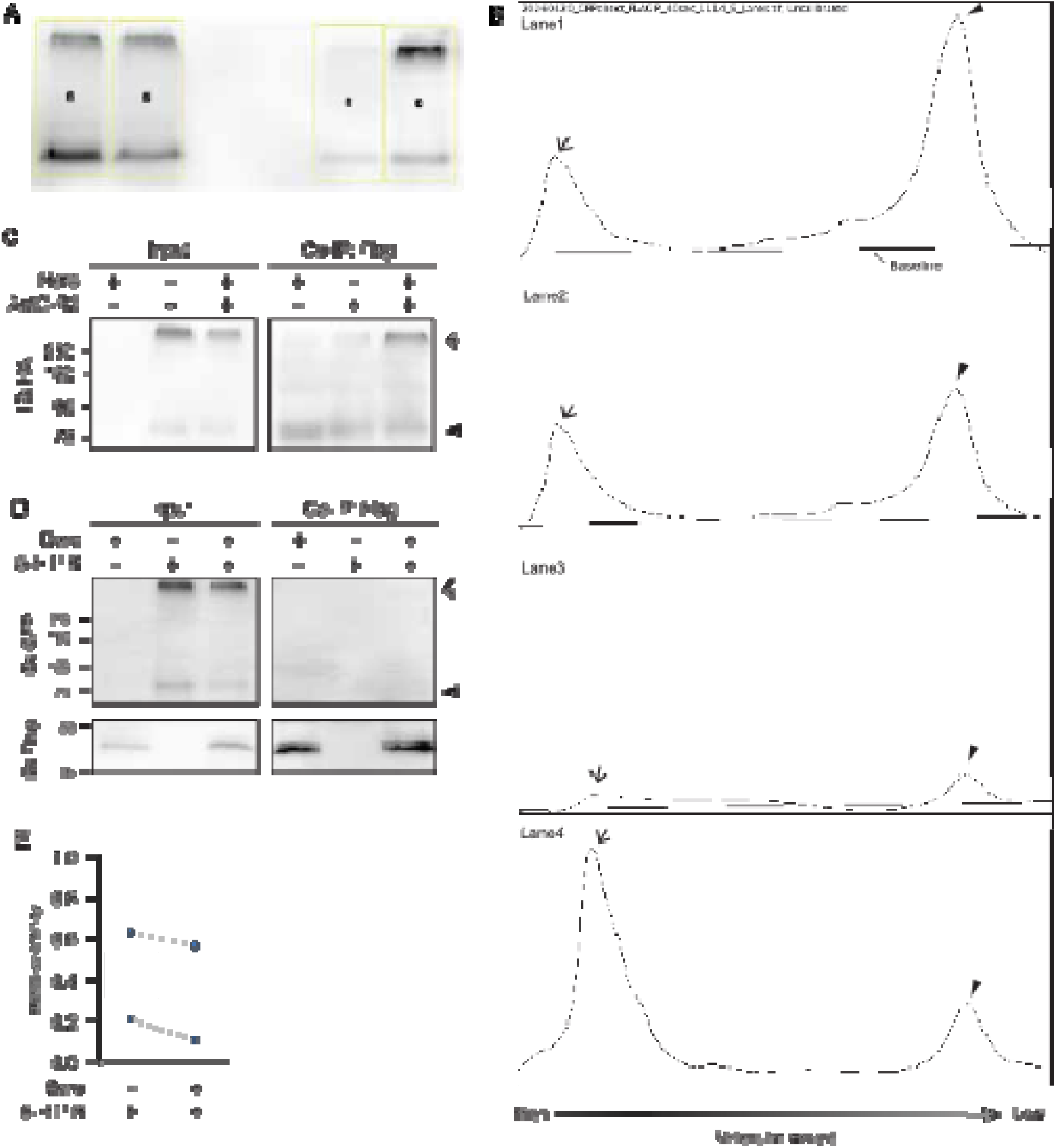
Co-immunoprecipitation analyses of Bero and GPCRs. (A) The Regions of interest (ROIs) defined using the Gels function in ImageJ for quantification of immunoblot signals. (B) Quantification of signals within the ROIs and determination of baselines. (C) Co-immunoprecipitation of Bero^FLAG^ with AstC-R2^HA&GFP^ in *Drosophila* S2 cells. Immunoblotting was performed using an anti-HA antibody. The triangle indicates putative monomeric AstC-R2, and the arrow indicates a high-molecular-weight AstC-R2 species. (D) Co-immunoprecipitation of Bero^FLAG^ with 5-HT1B^GFP^ in *Drosophila* S2 cells. Immunoblotting was performed using anti-GFP and anti-FLAG antibodies. The triangle indicates putative monomeric 5-HT1B, and the arrow indicates a high-molecular-weight 5-HT1B species. (E) Quantification of relative Bero–5-HT1B association.

**Figure S15.**
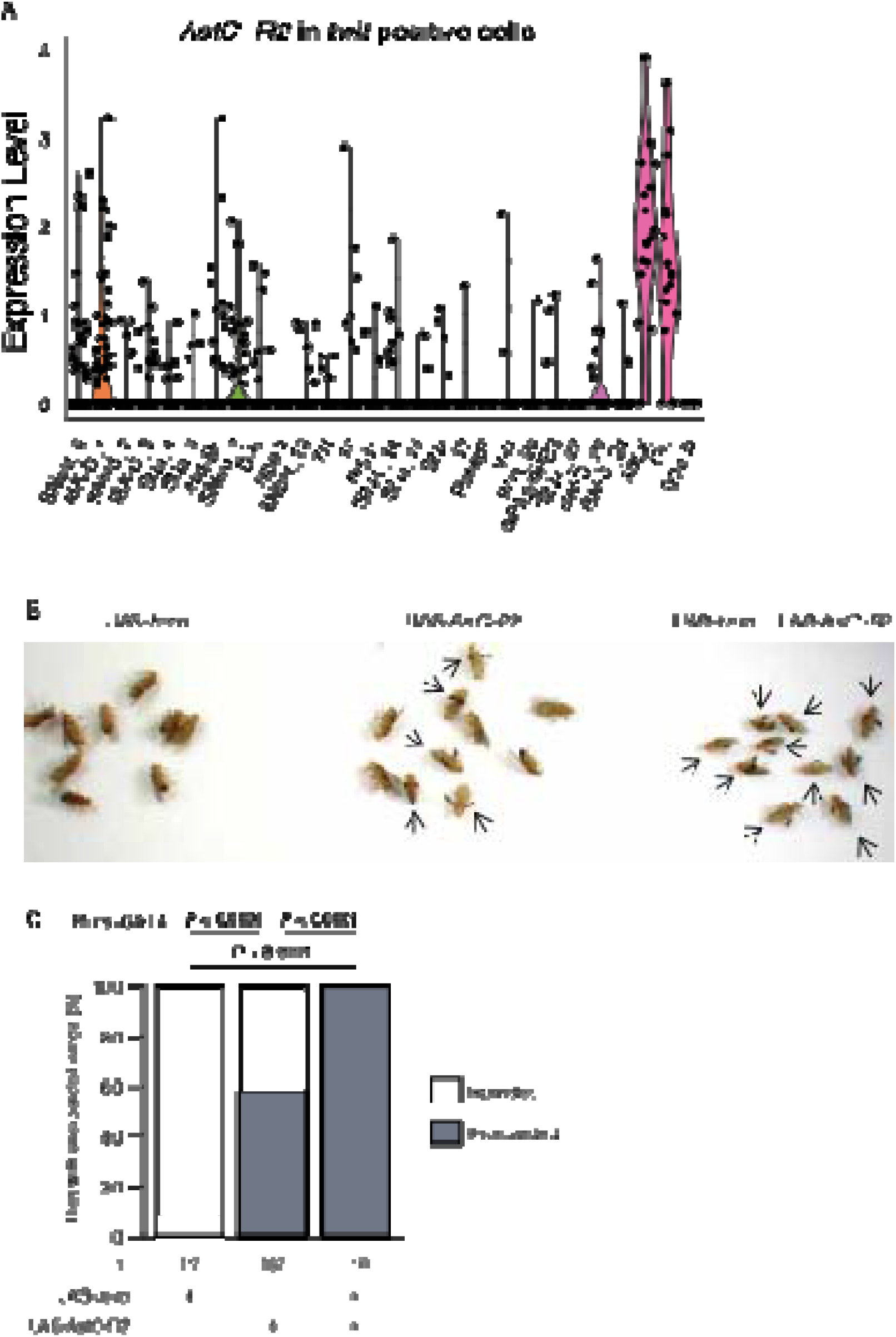
Ectopic expression of Bero and AstC-R2 in type III neuroendocrine cells. (A) Violin plot showing *AstC-R2* expression levels in a subset of *twit*-positive cells, which include motor neurons and neurosecretory cells. (B) Representative images of newly eclosed adult flies following ectopic expression of *bero*, *AstC-R2*, or both using *CCAP-GAL4*. Arrows indicate flies with unexpanded wings. (C) Proportion of adult flies with unexpanded wings following ectopic expression using the independent type III neuroendocrine cell driver *burs-GAL4*. Multiple comparisons with Fisher’s exact test.

**Figure S16.**
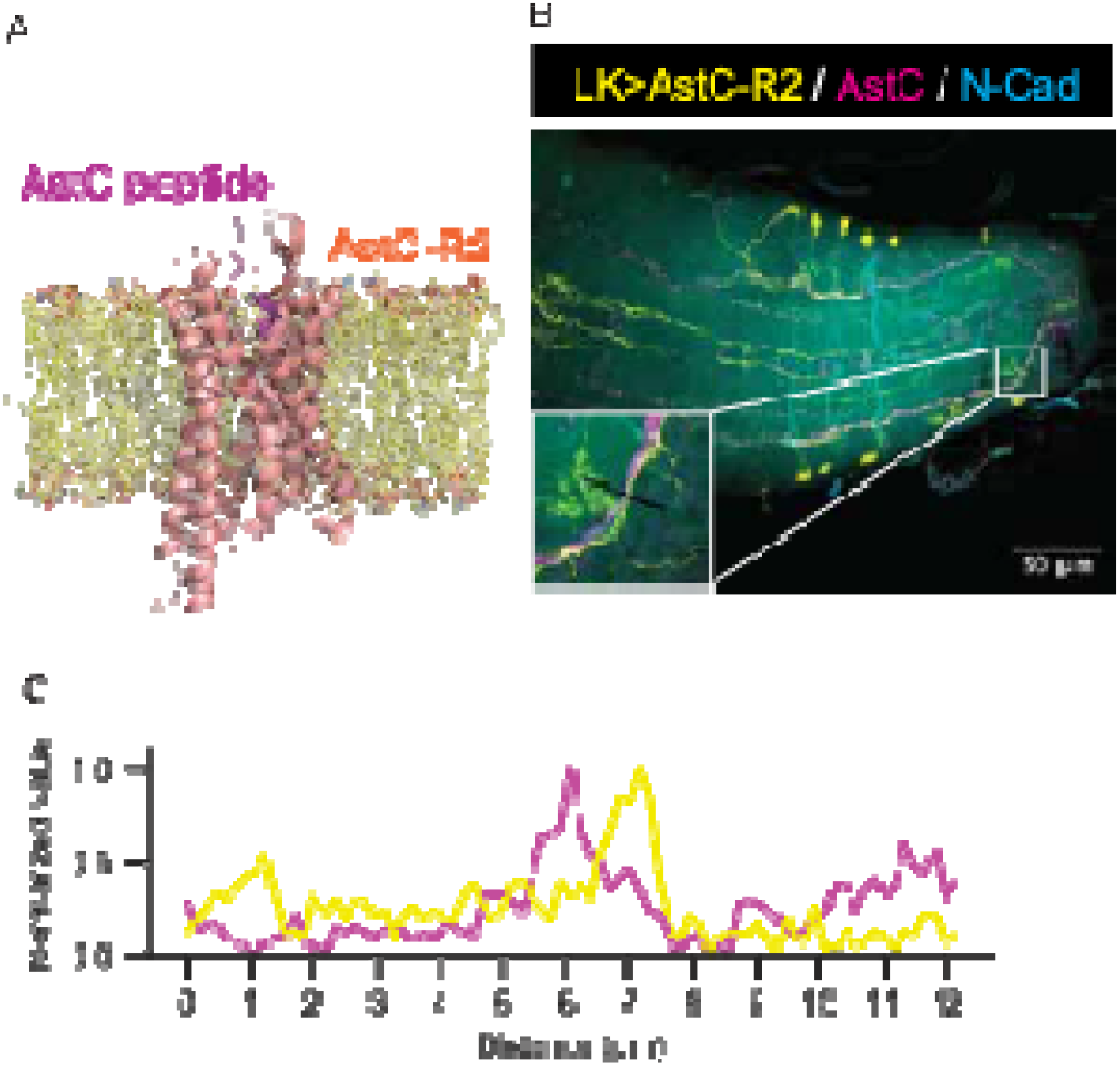
AstC^+^ neurites are in close proximity to AstC-R2-expressing neurites. (A) Predicted AstC–AstC-R2 complex structure generated using AlphaFold-Multimer and modeled in a membrane environment using CHARMM-GUI. (B) Confocal image showing AstC-R2::GFP expressed under *Lk-GAL4* (yellow), AstC immunoreactivity (magenta), and N-cadherin (cyan). Scale bar, 50 µm. (C) Normalized line profiles of AstC-R2 (yellow) and AstC (magenta) signals across the region indicated in (B).

**Figure S17.**
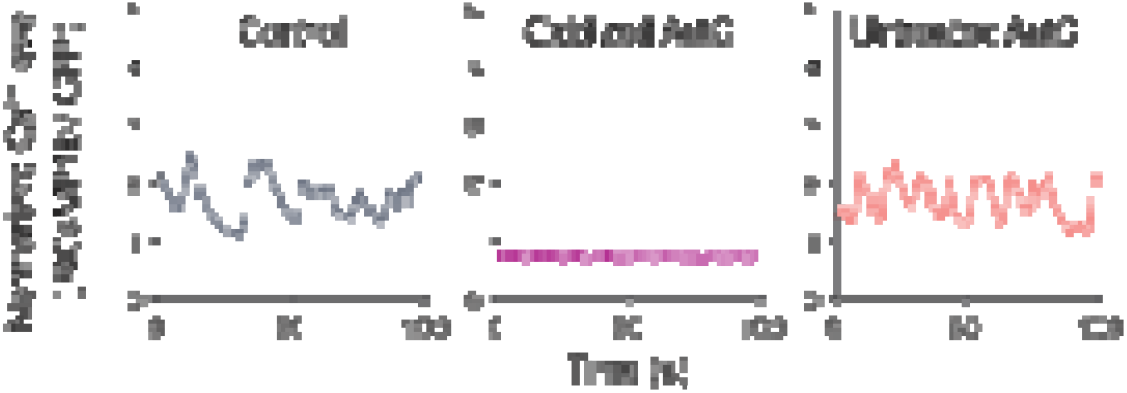
Oxidized AstC peptide decreases intracellular Ca²⁺ levels in ABLK neurons. Representative traces of normalized intracellular Ca²⁺ levels after treatment with oxidized or untreated AstC peptide.

**Figure S18.**
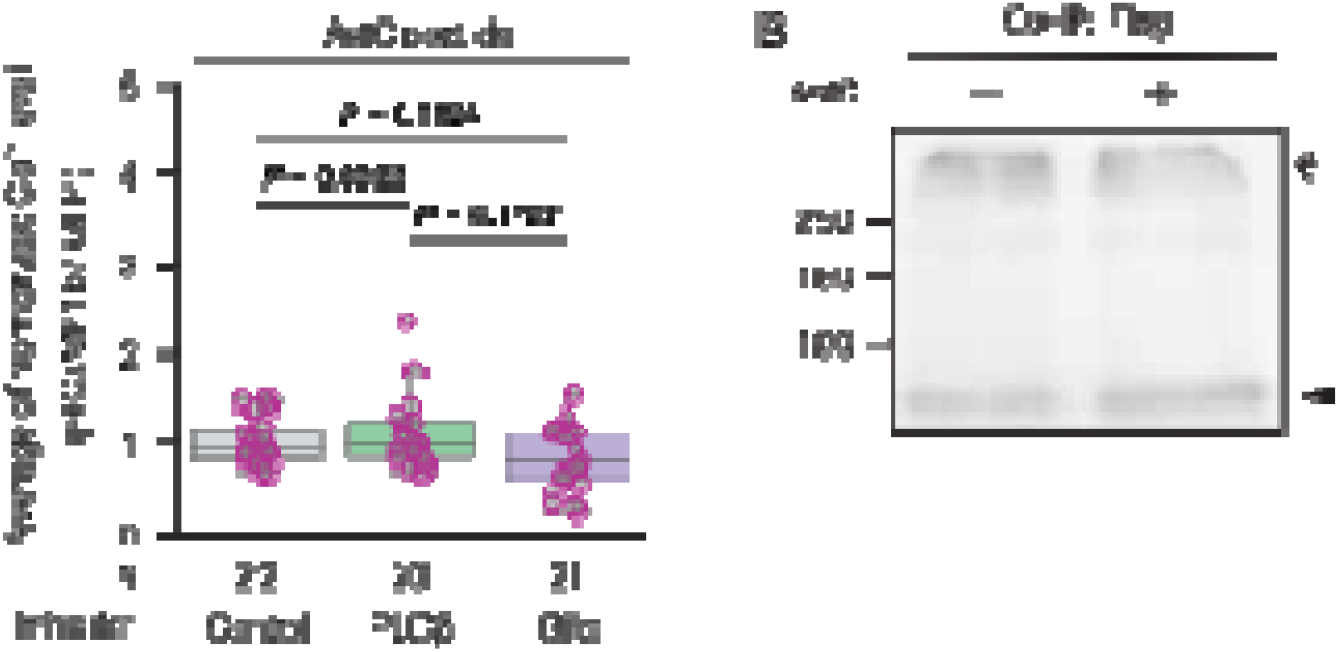
Effects of Gαi/o inhibition and AstC treatment on Bero–AstC-R2 signaling. (A) Quantification of intracellular Ca²⁺ levels in ABLK neurons following AstC treatment in the presence of the indicated inhibitors. Kruskal–Wallis test, followed by Dunn’s test. (B) Co-immunoprecipitation analysis of Bero and AstC-R2 in the absence or presence of AstC peptide.

**Figure S19.**
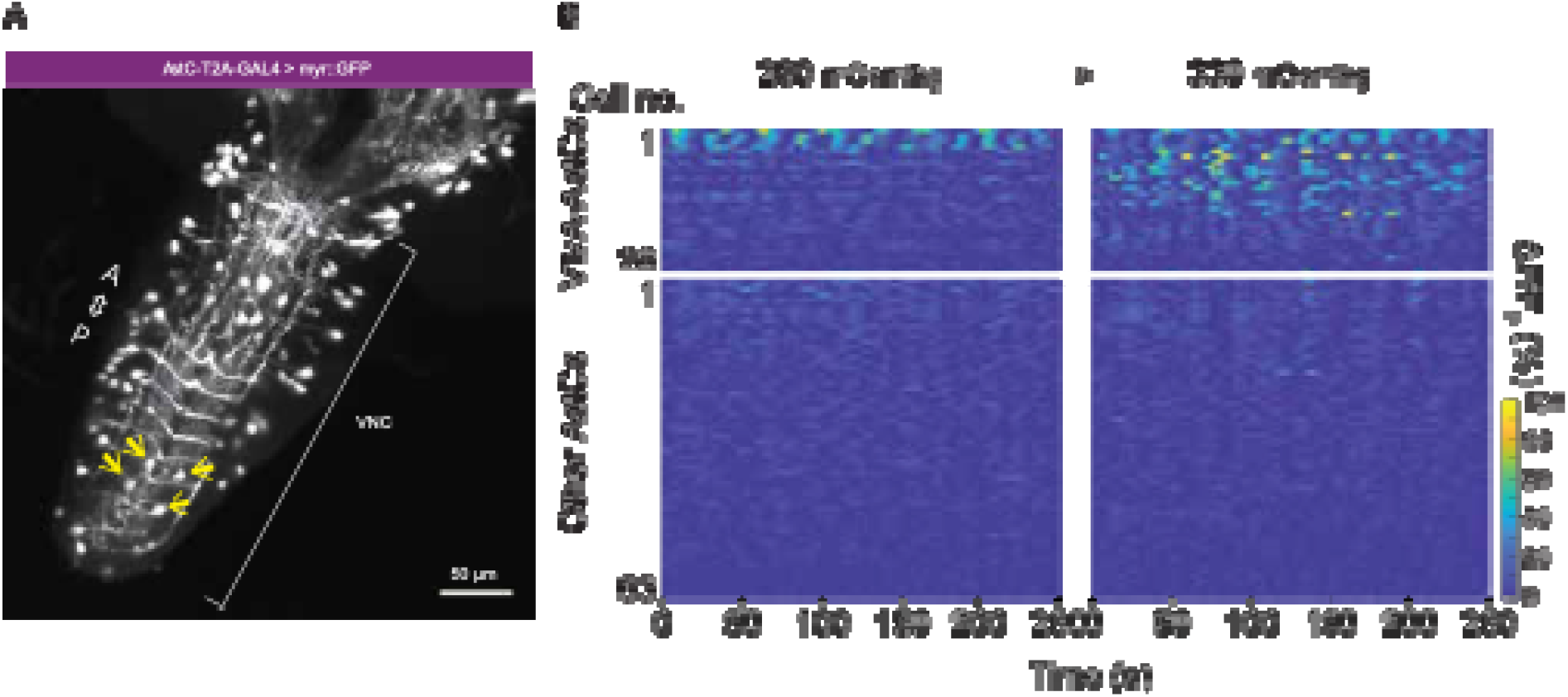
VMA-AstC neurons are osmosensitive. (A) Confocal image showing *AstC-T2A-GAL4*-labeled cells. Arrows indicate VMA-AstC neurons. Scale bar, 50 µm. (B) Representative heat maps showing spontaneous Ca²⁺ activity (ΔF/F₀) over time in AstC neurons in the VNC under 280 and 330 mOsm/kg conditions. Color intensity represents ΔF/F₀.

## SUPPLEMENTAL TABLES

**Table S1.**
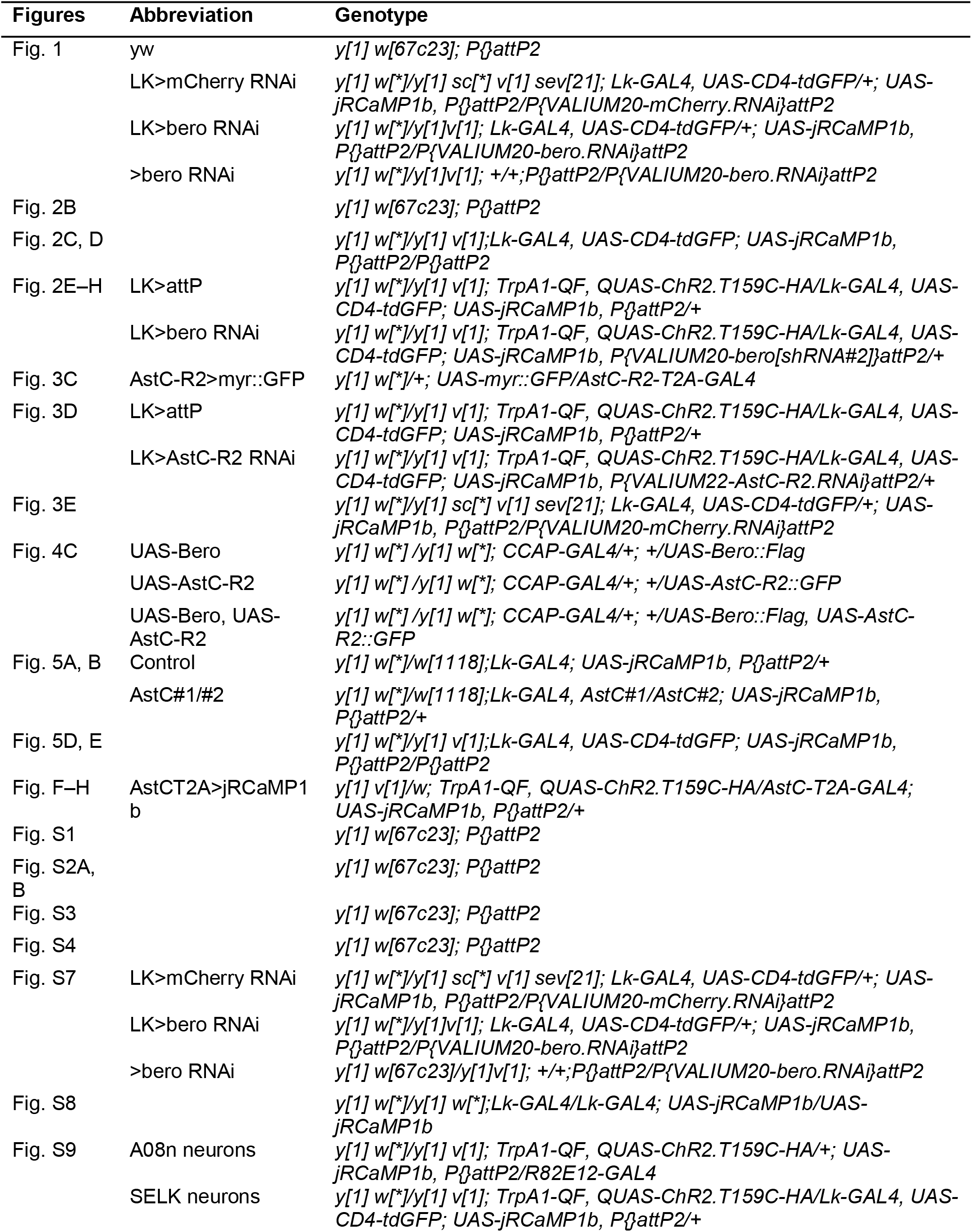
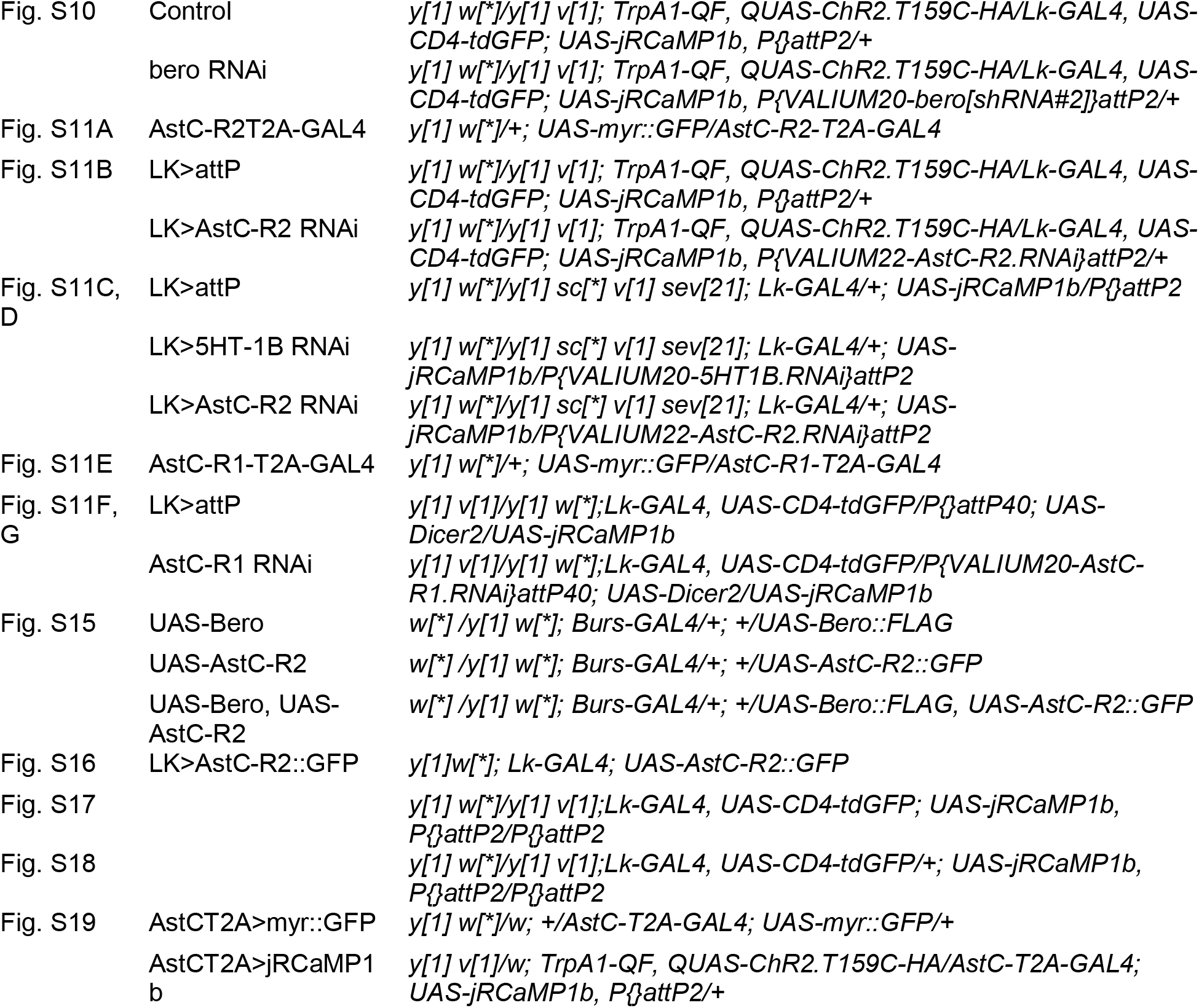
Fly genotypes used in this study Figures Abbreviation Genotype.

**Table S2.** Composition of saline solutions used in this study.

| **Saline solution** | **NaCl** | **KCl** | **CaCl<sub>2</sub>** | **MgCl<sub>2</sub>** | **NaHCO<sub>3</sub>** | **TES** | **HEPES** | **D-glucose** | **Sucrose** | **Trehalose** | **l-glucose** |
|---|---|---|---|---|---|---|---|---|---|---|---|
| S-280 | 120 | 3 | 1.5 | 4 | 10 | 5 | 10 | 10 | 0 | 0 | 0 |
| S-305 | 120 | 3 | 1.5 | 4 | 10 | 5 | 10 | 10 | 10 | 10 | 0 |
| S-330 | 120 | 3 | 1.5 | 4 | 10 | 5 | 10 | 10 | 20 | 20 | 0 |
| L-glucose, 330 mOsm/kg | 120 | 3 | 1.5 | 4 | 10 | 5 | 10 | 10 | 0 | 0 | 60 |

## REFERENCES

1. Flavell, S.W., Gogolla, N., Lovett-Barron, M., and Zelikowsky, M. (2022). The emergence and influence of internal states. Neuron 110, 2545–2570.

2. Jourjine, N. (2017). Hunger and thirst interact to regulate ingestive behavior in flies and mammals. Bioessays 39, 1600261.

3. Leib, D.E., Zimmerman, C.A., and Knight, Z.A. (2016). Thirst. Curr. Biol. 26, R1260–R1265.

4. Zimmerman, C.A., Lin, Y.-C., Leib, D.E., Guo, L., Huey, E.L., Daly, G.E., Chen, Y., and Knight, Z.A. (2016). Thirst neurons anticipate the homeostatic consequences of eating and drinking. Nature 537, 680–684.

5. Matsuda, T., Hiyama, T.Y., Niimura, F., Matsusaka, T., Fukamizu, A., Kobayashi, K., Kobayashi, K., and Noda, M. (2017). Distinct neural mechanisms for the control of thirst and salt appetite in the subfornical organ. Nat. Neurosci. 20, 230–241.

6. Matsuda, T., Hiyama, T.Y., Kobayashi, K., Kobayashi, K., and Noda, M. (2020). Distinct CCK-positive SFO neurons are involved in persistent or transient suppression of water intake. Nat. Commun. 11, 5692.

7. Zhang, F., Mak, S.O.K., Liu, Y., Ke, Y., Rao, F., Yung, W.H., Zhang, L., and Chow, B.K.C. (2022). Secretin receptor deletion in the subfornical organ attenuates the activation of excitatory neurons under dehydration. Curr. Biol. 32, 4832–4841.e5.

8. González-Segarra, A.J., Pontes, G., Jourjine, N., Toro, A.D., and Scott, K. (2023). Hunger-and thirst-sensing neurons modulate a neuroendocrine network to coordinate sugar and water ingestion. Elife 12, 1–24.

9. Senapati, B., Tsao, C.-H., Juan, Y.-A., Chiu, T.-H., Wu, C.-L., Waddell, S., and Lin, S. (2019). A neural mechanism for deprivation state-specific expression of relevant memories in Drosophila. Nat. Neurosci. 22, 2029–2039.

10. Chu, L.-A., Tai, C.-Y., and Chiang, A.-S. (2024). Thirst-driven hygrosensory suppression promotes water seeking in Drosophila. Proc. Natl. Acad. Sci. U. S. A. 121, 2017.

11. Johnson, W.A., and Carder, J.W. (2012). Drosophila nociceptors mediate larval aversion to dry surface environments utilizing both the painless TRP Channel and the DEG/ENaC subunit, PPK1. PLoS One 7. 10.1371/JOURNAL.PONE.0032878.

12. Rodriguez, L., Sokolowski, M.B., and Shore, J.S. (1992). Habitat selection by *Drosophila melanogaster* larvae. J. Evol. Biol. 5, 61–70.

13. Kanaoka, Y., Onodera, K., Watanabe, K., Hayashi, Y., Usui, T., Uemura, T., and Hattori, Y. (2023). Inter-organ Wingless/Ror/Akt signaling regulates nutrient-dependent hyperarborization of somatosensory neurons. Elife 12. 10.7554/eLife.79461.

14. Lampe, L., Newell, C.L., Wang, B.-J., Bailey, A.P., Makki, R., Alexandre, C., Gilmore, I.S., Zhao, L., and Gould, A.P. (2026). Identification of a specialized lipid barrier for *Drosophila* metamorphosis. Sci. Adv. 12. 10.1126/sciadv.adw6168.

15. Vogt, K., Zimmerman, D.M., Schlichting, M., Hernandez-Nunez, L., Qin, S., Malacon, K., Rosbash, M., Pehlevan, C., Cardona, A., and Samuel, A.D.T. (2021). Internal state configures olfactory behavior and early sensory processing in Drosophila larvae. Sci. Adv. 7, eabd6900.

16. Kim, S.M., Su, C.-Y., and Wang, J.W. (2017). Neuromodulation of innate behaviors in Drosophila. Annu. Rev. Neurosci. 40, 327–348.

17. Soden, M.E., Yee, J.X., and Zweifel, L.S. (2023). Circuit coordination of opposing neuropeptide and neurotransmitter signals. Nature 619, 332–337.

18. Grove, J.C.R., Gray, L.A., La Santa Medina, N., Sivakumar, N., Ahn, J.S., Corpuz, T.V., Berke, J.D., Kreitzer, A.C., and Knight, Z.A. (2022). Dopamine subsystems that track internal states. Nature 608, 374–380.

19. Boivin, J.-C., Zhao, Y.Q., Zhu, J., Dakin, J.T., Ning, J., and Ohyama, T. (2026). A positive feedback loop between sensory and octopaminergic neurons underlies nociceptive plasticity in Drosophila larvae. PLoS Genet. 22, e1012122.

20. Loughner, C.L., Bruford, E.A., McAndrews, M.S., Delp, E.E., Swamynathan, S., and Swamynathan, S.K. (2016). Organization, evolution and functions of the human and mouse Ly6/uPAR family genes. Hum. Genomics 10, 10.

21. Zaigraev, M.M., Lyukmanova, E.N., Paramonov, A.S., Shenkarev, Z.O., and Chugunov, A.O. (2022). Orientational Preferences of GPI-Anchored Ly6/uPAR Proteins. Int. J. Mol. Sci. 24, 11.

22. Lyukmanova, E.N., Shenkarev, Z.O., Shulepko, M.A., Mineev, K.S., D’Hoedt, D., Kasheverov, I.E., Filkin, S.Y., Krivolapova, A.P., Janickova, H., Dolezal, V., et al. (2011). NMR structure and action on nicotinic acetylcholine receptors of water-soluble domain of human LYNX1. J. Biol. Chem. 286, 10618–10627.

23. Morishita, H., Miwa, J.M., Heintz, N., and Hensch, T.K. (2010). Lynx1, a Cholinergic Brake, Limits Plasticity in Adult Visual Cortex. Science 330, 1238–1240.

24. Koh, K., Joiner, W.J., Wu, M.N., Yue, Z., Smith, C.J., and Sehgal, A. (2008). Identification of SLEEPLESS, a Sleep-Promoting Factor. Science 321, 372–376.

25. Wu, M.N., Joiner, W.J., Dean, T., Yue, Z., Smith, C.J., Chen, D., Hoshi, T., Sehgal, A., and Koh, K. (2010). SLEEPLESS, a Ly-6/neurotoxin family member, regulates the levels, localization and activity of Shaker. Nat. Neurosci. 13, 69–75.

26. Wu, M., Robinson, J.E., and Joiner, W.J. (2014). SLEEPLESS Is a Bifunctional Regulator of Excitability and Cholinergic Synaptic Transmission. Curr. Biol. 24, 621–629.

27. Wu, M., Puddifoot, C.A., Taylor, P., and Joiner, W.J. (2015). Mechanisms of Inhibition and Potentiation of α4β2 Nicotinic Acetylcholine Receptors by Members of the Ly6 Protein Family. J. Biol. Chem. 290, 24509–24518.

28. Özhan, G., Sezgin, E., Wehner, D., Pfister, A.S., Kühl, S.J., Kagermeier-Schenk, B., Kühl, M., Schwille, P., and Weidinger, G. (2013). Lypd6 enhances wnt/β-catenin signaling by promoting lrp6 phosphorylation in raft plasma Membrane Domains. Dev. Cell 26, 331–345.

29. Resnati, M., Pallavicini, I., Wang, J.M., Oppenheim, J., Serhan, C.N., Romano, M., and Blasi, F. (2002). The fibrinolytic receptor for urokinase activates the G protein-coupled chemotactic receptor FPRL1/LXA4R. Proc. Natl. Acad. Sci. U. S. A. 99, 1359–1364.

30. Gorrasi, A., Li Santi, A., Amodio, G., Alfano, D., Remondelli, P., Montuori, N., and Ragno, P. (2014). The urokinase receptor takes control of cell migration by recruiting integrins and FPR1 on the cell surface. PLoS One 9, e86352.

31. Minopoli, M., Polo, A., Ragone, C., Ingangi, V., Ciliberto, G., Pessi, A., Sarno, S., Budillon, A., Costantini, S., and Carriero, M.V. (2019). Structure-function relationship of an Urokinase Receptor-derived peptide which inhibits the Formyl Peptide Receptor type 1 activity. Sci. Rep. 9, 12169.

32. Gorrasi, A., Petrone, A.M., Li Santi, A., Alfieri, M., Montuori, N., and Ragno, P. (2020). New pieces in the puzzle of uPAR role in cell migration mechanisms. Cells 9, E2531.

33. Li, K., Tsukasa, Y., Kurio, M., Maeta, K., Tsumadori, A., Baba, S., Nishimura, R., Murakami, A., Onodera, K., Morimoto, T., et al. (2023). Belly roll, a GPI-anchored Ly6 protein, regulates Drosophila melanogaster escape behaviors by modulating the excitability of nociceptive peptidergic interneurons. Elife 12, 1–31.

34. Tsukasa, Y., Uemura, T., and Usui, T. (2026). A GPI-anchored Ly6/ uPAR superfamily gene *belly roll* is expressed in multiple peptidergic neurons in *Drosophila melanogaster* larvae. Genes Cells 31. 10.1111/gtc.70116.

35. Imambocus, B.N., Zhou, F., Formozov, A., Wittich, A., Tenedini, F.M., Hu, C., Sauter, K., Macarenhas Varela, E., Herédia, F., Casimiro, A.P., et al. (2022). A neuropeptidergic circuit gates selective escape behavior of Drosophila larvae. Curr. Biol. 32, 149–163.e8.

36. Hu, Y., Wang, C., Yang, L., Pan, G., Liu, H., Yu, G., and Ye, B. (2020). A Neural Basis for Categorizing Sensory Stimuli to Enhance Decision Accuracy. Curr. Biol. 30, 4896–4909.e6.

37. Okusawa, S., Kohsaka, H., and Nose, A. (2014). Serotonin and Downstream Leucokinin Neurons Modulate Larval Turning Behavior in Drosophila. J. Neurosci. 34, 2544–2558.

38. Benoit, J.B., McCluney, K.E., Degennaro, M.J., and Dow, J.A.T. (2023). Dehydration Dynamics in Terrestrial Arthropods: From Water Sensing to Trophic Interactions. Annu. Rev. Entomol. 68, 129– 149.

39. Halberg, K.V., and Denholm, B. (2024). Mechanisms of Systemic Osmoregulation in Insects. Annu. Rev. Entomol. 69, 415–438.

40. Fanara, J.J., Sassi, P.L., Goenaga, J., and Hasson, E. (2024). Genetic basis and repeatability for desiccation resistance in Drosophila melanogaster (Diptera: Drosophilidae). Genetica 152, 1–9.

41. Thorat, L.J., Gaikwad, S.M., and Nath, B.B. (2012). Trehalose as an indicator of desiccation stress in Drosophila melanogaster larvae: A potential marker of anhydrobiosis. Biochem. Biophys. Res. Commun. 419, 638–642.

42. Kawano, T., Shimoda, M., Matsumoto, H., Ryuda, M., Tsuzuki, S., and Hayakawa, Y. (2010). Identification of a Gene, Desiccate, Contributing to Desiccation Resistance in Drosophila melanogaster. J. Biol. Chem. 285, 38889–38897.

43. Wang, Z., Receveur, J.P., Pu, J., Cong, H., Richards, C., Liang, M., and Chung, H. (2022). Desiccation resistance differences in Drosophila species can be largely explained by variations in cuticular hydrocarbons. Elife 11, 1–22.

44. Casares, P., Carracedo, M.C., and García-Florez, L. (1997). Analysis of larval behaviours underlying the pupation height phenotype in Drosophila simulans and D melanogaster. Genet. Sel. Evol. 29, 589–600.

45. Sokolowski, M.B. (1985). Genetics and ecology of Drosophila melanogaster larval foraging and pupation behaviour. J. Insect Physiol. 31, 857–864.

46. Sokolowski, M.B., Bauer, S.J., Wai-Ping, V., Rodriguez, L., Wong, J.L., and Kent, C. (1986). Ecological genetics and behaviour of Drosophila melanogaster larvae in nature. Anim. Behav. 34, 403–408.

47. Yamanaka, N., Romero, N.M., Martin, F.A., Rewitz, K.F., Sun, M., O’Connor, M.B., and Léopold, P. (2013). Neuroendocrine Control of Drosophila Larval Light Preference. Science 341, 1113–1116.

48. Turner, H.N., Armengol, K., Patel, A.A., Himmel, N.J., Sullivan, L., Iyer, S.C., Bhattacharya, S., Iyer, E.P.R., Landry, C., Galko, M.J., et al. (2016). The TRP Channels Pkd2, NompC, and Trpm Act in Cold-Sensing Neurons to Mediate Unique Aversive Behaviors to Noxious Cold in Drosophila. Curr. Biol. 26, 3116–3128.

49. Jovanic, T., Schneider-Mizell, C.M., Shao, M., Masson, J.-B., Denisov, G., Fetter, R.D., Mensh, B.D., Truman, J.W., Cardona, A., and Zlatic, M. (2016). Competitive Disinhibition Mediates Behavioral Choice and Sequences in Drosophila. Cell 167, 858–870.e19.

50. Masson, J.-B., Laurent, F., Cardona, A., Barré, C., Skatchkovsky, N., Zlatic, M., and Jovanic, T. (2020). Identifying neural substrates of competitive interactions and sequence transitions during mechanosensory responses in Drosophila. PLoS Genet. 16, e1008589.

51. Tracey, W.D., Wilson, R.I., Laurent, G., and Benzer, S. (2003). painless, a Drosophila Gene Essential for Nociception. Cell 113, 261–273.

52. Hwang, R.Y., Zhong, L., Xu, Y., Johnson, T., Zhang, F., Deisseroth, K., and Tracey, W.D. (2007). Nociceptive Neurons Protect Drosophila Larvae from Parasitoid Wasps. Curr. Biol. 17, 2105–2116.

53. Terada, S.-I., Matsubara, D., Onodera, K., Matsuzaki, M., Uemura, T., and Usui, T. (2016). Neuronal processing of noxious thermal stimuli mediated by dendritic Ca2+ influx in Drosophila somatosensory neurons. Elife 5. 10.7554/eLife.12959.

54. Onodera, K., Baba, S., Murakami, A., Uemura, T., and Usui, T. (2017). Small conductance Ca2+-activated K+ channels induce the firing pause periods during the activation of Drosophila nociceptive neurons. Elife 6. 10.7554/eLife.29754.

55. Xiang, Y., Yuan, Q., Vogt, N., Looger, L.L., Jan, L.Y., and Jan, Y.N. (2010). Light-avoidance-mediating photoreceptors tile the Drosophila larval body wall. Nature 468, 921–926.

56. Ohyama, T., Schneider-Mizell, C.M., Fetter, R.D., Aleman, J.V., Franconville, R., Rivera-Alba, M., Mensh, B.D., Branson, K.M., Simpson, J.H., Truman, J.W., et al. (2015). A multilevel multimodal circuit enhances action selection in Drosophila. Nature 520, 633–639.

57. Grueber, W.B., Jan, L.Y., and Jan, Y.N. (2002). Tiling of the Drosophila epidermis by multidendritic sensory neurons. Development 129, 2867–2878.

58. Imambocus, B.N., Jiang, Y., Schleyer, M., and Soba, P. (2025). Peptidergic top-down control of metabolic state-dependent behavioral decisions in a conflicting sensory context. bioRxiv, 2025.03.12.642538. 10.1101/2025.03.12.642538.

59. Boivin, J.-C., Zhu, J., and Ohyama, T. (2023). Nociception in fruit fly larvae. Front. Pain Res. (Lausanne) 4. 10.3389/fpain.2023.1076017.

60. Zandawala, M., Marley, R., Davies, S.A., and Nässel, D.R. (2018). Characterization of a set of abdominal neuroendocrine cells that regulate stress physiology using colocalized diuretic peptides in Drosophila. Cell. Mol. Life Sci. 75, 1099–1115.

61. Nässel, D.R. (2021). Leucokinin and Associated Neuropeptides Regulate Multiple Aspects of Physiology and Behavior in Drosophila. Int. J. Mol. Sci. 22, 1940.

62. Jourjine, N., Mullaney, B.C., Mann, K., and Scott, K. (2016). Coupled Sensing of Hunger and Thirst Signals Balances Sugar and Water Consumption. Cell 166, 855–866.

63. Pool, A.H., Wang, T., Stafford, D.A., Chance, R.K., Lee, S., Ngai, J., and Oka, Y. (2020). The cellular basis of distinct thirst modalities. Nature 588. 10.1038/s41586-020-2821-8.

64. Kurio, M., Tsukasa, Y., Uemura, T., and Usui, T. (2024). Refinement of a technique for collecting and evaluating the osmolality of haemolymph from Drosophila larvae. J. Exp. Biol. 227, 1–6.

65. Fernando Vieira Contreras, A., Auger, G.M., Ljaschenko, D., Blanco-Redondo, B., and Langenhan Correspondence, T. (2024). The adhesion G-protein-coupled receptor mayo/ CG11318 controls midgut development in Drosophila. CellReports 43, 113640.

66. Chen, L., Liu, C., and Liu, L. (2009). Osmolality-induced tuning of action potentials in trigeminal ganglion neurons. Neurosci. Lett. 452, 79–83.

67. Allen, W.E., Chen, M.Z., Pichamoorthy, N., Tien, R.H., Pachitariu, M., Luo, L., and Deisseroth, K. (2019). Thirst regulates motivated behavior through modulation of brainwide neural population dynamics. Science 364. 10.1126/science.aav3932.

68. Hu, C., Petersen, M., Hoyer, N., Spitzweck, B., Tenedini, F., Wang, D., Gruschka, A., Burchardt, L.S., Szpotowicz, E., Schweizer, M., et al. (2017). Sensory integration and neuromodulatory feedback facilitate Drosophila mechanonociceptive behavior. Nat. Neurosci. 20, 1085–1095.

69. Dillon, N., Cocanougher, B., Sood, C., Yuan, X., Kohn, A.B., Moroz, L.L., Siegrist, S.E., Zlatic, M., and Doe, C.Q. (2022). Single cell RNA-seq analysis reveals temporally-regulated and quiescence-regulated gene expression in Drosophila larval neuroblasts. Neural Dev. 17, 7.

70. Corrales, M., Cocanougher, B.T., Kohn, A.B., Wittenbach, J.D., Long, X.S., Lemire, A., Cardona, A., Singer, R.H., Moroz, L.L., and Zlatic, M. (2022). A single-cell transcriptomic atlas of complete insect nervous systems across multiple life stages. Neural Dev. 17, 8.

71. Nguyen, T.H., Vicidomini, R., Choudhury, S.D., Han, T.H., Maric, D., Brody, T., and Serpe, M. (2024). scRNA-seq data from the larval Drosophila ventral cord provides a resource for studying motor systems function and development. Dev. Cell 59, 1210–1230.e9.

72. Kondo, S., Takahashi, T., Yamagata, N., Imanishi, Y., Katow, H., Hiramatsu, S., Lynn, K., Abe, A., Kumaraswamy, A., and Tanimoto, H. (2020). Neurochemical Organization of the Drosophila Brain Visualized by Endogenously Tagged Neurotransmitter Receptors. Cell Rep. 30, 284–297.e5.

73. Işbilir, A., Duan Sahbaz, B., Tuncgenc, G., Bünemann, M., Lohse, M.J., and Birgül-Iyison, N. (2020). Pharmacological Characterization of the Stick Insect Carausius morosus Allatostatin-C Receptor with Its Endogenous Agonist. ACS Omega 5, 32183–32194.

74. Shahraki, A., Işbilir, A., Dogan, B., Lohse, M.J., Durdagi, S., and Birgul-Iyison, N. (2021). Structural and Functional Characterization of Allatostatin Receptor Type-C of Thaumetopoea pityocampa , a Potential Target for Next-Generation Pest Control Agents. J. Chem. Inf. Model. 61, 715–728.

75. Kahveci, K., Düzgün, M.B., Atis, A.E., Yılmaz, A., Shahraki, A., Coskun, B., Durdagi, S., and Birgul Iyison, N. (2024). Discovering allatostatin type-C receptor specific agonists. Nat. Commun. 15, 3965.

76. Urlacher, E., Soustelle, L., Parmentier, M.L., Verlinden, H., Gherardi, M.J., Fourmy, D., Mercer, A.R., Devaud, J.M., and Massou, I. (2016). Honey bee allatostatins target galanin/somatostatin-like receptors and modulate learning: A conserved function? PLoS One 11. 10.1371/journal.pone.0146248.

77. Robertson, M.J., Meyerowitz, J.G., Panova, O., Borrelli, K., and Skiniotis, G. (2022). Plasticity in ligand recognition at somatostatin receptors. Nat. Struct. Mol. Biol. 29, 210–217.

78. Sgourakis, N.G., Bagos, P.G., Papasaikas, P.K., and Hamodrakas, S.J. (2005). A method for the prediction of GPCRs coupling specificity to G-proteins using refined profile Hidden Markov Models. BMC Bioinformatics 6. 10.1186/1471-2105-6-104.

79. Pfeil, E.M., Brands, J., Merten, N., Vögtle, T., Vescovo, M., Rick, U., Albrecht, I.M., Heycke, N., Kawakami, K., Ono, Y., et al. (2020). Heterotrimeric G Protein Subunit Gαq Is a Master Switch for Gβγ-Mediated Calcium Mobilization by Gi-Coupled GPCRs. Mol. Cell 80, 940–954.e6.

80. Evans, R., O’neill, M., Pritzel, A., Antropova, N., Senior, A., Green, T., Žídek, A., Bates, R., Blackwell, S., Yim, J., et al. (2021). Protein complex prediction with AlphaFold-Multimer. bioRxivorg. 10.1101/2021.10.04.463034.

81. Mirdita, M., Schütze, K., Moriwaki, Y., Heo, L., Ovchinnikov, S., and Steinegger, M. (2022). ColabFold: making protein folding accessible to all. Nat. Methods 19, 679–682.

82. Holze, J., Bermudez, M., Pfeil, E.M., Kauk, M., Bödefeld, T., Irmen, M., Matera, C., Dallanoce, C., De Amici, M., Holzgrabe, U., et al. (2020). Ligand-Specific Allosteric Coupling Controls G-Protein-Coupled Receptor Signaling. ACS Pharmacol. Transl. Sci. 3, 859–867.

83. Chen, S., Teng, X., and Zheng, S. (2023). Molecular basis for the selective G protein signaling of somatostatin receptors. Nat. Chem. Biol. 19, 133–140.

84. Pfeiffer, M., Koch, T., Schröder, H., Klutzny, M., Kirscht, S., Kreienkamp, H.J., Höllt, V., and Schulz, S. (2001). Homo- and heterodimerization of somatostatin receptor subtypes. J. Biol. Chem. 276, 14027–14036.

85. Perret, B.G., Wagner, R., Lecat, S., Brillet, K., Rabut, G., Bucher, B., and Pattus, F. (2003). Expression of EGFP-amino-tagged human mu opioid receptor in Drosophila Schneider 2 cells: a potential expression system for large-scale production of G-protein coupled receptors. Protein Expr. Purif. 31, 123–132.

86. Luan, H., Peabody, N.C., Vinson, C.R., and White, B.H. (2006). Refined spatial manipulation of neuronal function by combinatorial restriction of transgene expression. Neuron 52, 425–436.

87. Tabuchi, M., Monaco, J.D., Duan, G., Bell, B., Liu, S., Liu, Q., Zhang, K., and Wu, M.N. (2018). Clock-Generated Temporal Codes Determine Synaptic Plasticity to Control Sleep. Cell 175, 1213–1227.e18.

88. Kreienkamp, H.J., Larusson, H.J., Witte, I., Roeder, T., Birgül, N., Hönck, H.H., Harder, S., Ellinghausen, G., Buck, F., and Richter, D. (2002). Functional annotation of two orphan G-protein-coupled receptors, drostar1 and -2, from Drosophila melanogaster and their ligands by reverse pharmacology. J. Biol. Chem. 277, 39937–39943.

89. Williamson, M., Lenz, C., Winther, Å.M.E., Nässel, D.R., and Grimmelikhuijzen, C.J.P. (2001). Molecular Cloning, Genomic Organization, and Expression of a C-Type (Manduca sexta-Type) Allatostatin Preprohormone from Drosophila melanogaster. Biochem. Biophys. Res. Commun. 282, 124–130.

90. Díaz, M.M., Schlichting, M., Abruzzi, K.C., Long, X., and Rosbash, M. (2019). Allatostatin-C/AstC-R2 Is a Novel Pathway to Modulate the Circadian Activity Pattern in Drosophila. Curr. Biol. 29, 13–22.e3.

91. Liu, J., Liu, W., Thakur, D., Mack, J., Spina, A., and Montell, C. (2023). Alleviation of thermal nociception depends on heat-sensitive neurons and a TRP channel in the brain. Curr. Biol. 33. 10.1016/j.cub.2023.04.055.

92. Zhang, Y., Yañez-Guerra, L.A., Tinoco, A.B., Castelán, N.E., Egertová, M., and Elphick, M.R. (2022). Somatostatin-type and allatostatin-C-type neuropeptides are paralogous and have opposing myoregulatory roles in an echinoderm. Proc. Natl. Acad. Sci. U. S. A. 119. 10.1073/pnas.2113589119.

93. Zeng, X., Komanome, Y., Kawasaki, T., Inada, K., Jonaitis, J., Pulver, S.R., Kazama, H., and Nose, A. (2021). An electrically coupled pioneer circuit enables motor development via proprioceptive feedback in Drosophila embryos. Curr. Biol. 31, 5327–5340.e5.

94. Meiselman, M.R., Alpert, M.H., Cui, X., Shea, J., Gregg, I., Gallio, M., and Yapici, N. (2022). Recovery from cold-induced reproductive dormancy is regulated by temperature-dependent AstC signaling. Curr. Biol. 32, 1362–1375.e8.

95. Price, M. (2002). Drosophila melanogaster flatline encodes a myotropin orthologue to Manduca sexta allatostatin. Peptides 23, 787–794.

96. Nissen, N.I., Anderson, K.R., Wang, H., Lee, H.S., Garrison, C., Eichelberger, S.A., Ackerman, K., Im, W., and Miwa, J.M. (2018). Augmenting the antinociceptive effects of nicotinic acetylcholine receptor activity through lynx1 modulation. PLoS One 13, e0199643.

97. Wu, M., Liu, C.Z., and Joiner, W.J. (2016). Structural Analysis and Deletion Mutagenesis Define Regions of QUIVER/SLEEPLESS that Are Responsible for Interactions with Shaker-Type Potassium Channels and Nicotinic Acetylcholine Receptors. PLoS One 11, e0148215.

98. Farrell, S.R., Rankin, D.R., Brecha, N.C., and Barnes, S. (2014). Somatostatin receptor subtype 4 modulates L-type calcium channels via Gβγ and PKC signaling in rat retinal ganglion cells. Channels (Austin) 8, 519–527.

99. Tomura, H., Okajima, F., Akbar, M., Abdul Majid, M., Sho, K., and Kondo, Y. (1994). Transfected human somatostatin receptor type 2, SSTR2, not only inhibits adenylate cyclase but also stimulates phospholipase C and Ca2+ mobilization. Biochem. Biophys. Res. Commun. 200, 986–992.

100. Ohashi, H., and Sakai, T. (2018). Leucokinin signaling regulates hunger–driven reduction of behavioral responses to noxious heat in Drosophila. Biochem. Biophys. Res. Commun. 499, 221– 226.

101. Veenstra, J.A., Agricola, H.-J., and Sellami, A. (2008). Regulatory peptides in fruit fly midgut. Cell Tissue Res. 334, 499–516.

102. Picao-Osorio, J., Johnston, J., Landgraf, M., Berni, J., and Alonso, C.R. (2015). MicroRNA-encoded behavior in Drosophila. Science 350, 815–820.

103. Dhar, G., Mukherjee, S., Nayak, N., Sahu, S., Bag, J., Rout, R., and Mishra, M. (2020). Various behavioural assays to detect the neuronal abnormality in flies. In Springer Protocols Handbooks (Springer US), pp. 223–251.

104. Risse, B., Berh, D., Otto, N., Klämbt, C., and Jiang, X. (2017). FIMTrack: An open source tracking and locomotion analysis software for small animals. PLoS Comput. Biol. 13, e1005530.

105. Dana, H., Mohar, B., Sun, Y., Narayan, S., Gordus, A., Hasseman, J.P., Tsegaye, G., Holt, G.T., Hu, A., Walpita, D., et al. (2016). Sensitive red protein calcium indicators for imaging neural activity. Elife 5, e12727.

106. Berndt, A., Schoenenberger, P., Mattis, J., Tye, K.M., Deisseroth, K., Hegemann, P., and Oertner, T.G. (2011). High-efficiency channelrhodopsins for fast neuronal stimulation at low light levels. Proc. Natl. Acad. Sci. U. S. A. 108, 7595–7600.

107. Vecsey, C.G., Pírez, N., and Griffith, L.C. (2014). The Drosophila neuropeptides PDF and sNPF have opposing electrophysiological and molecular effects on central neurons. J. Neurophysiol. 111, 1033– 1045.

108. Oh, Y., Lai, J.S.-Y., Mills, H.J., Erdjument-Bromage, H., Giammarinaro, B., Saadipour, K., Wang, J.G., Abu, F., Neubert, T.A., and Suh, G.S.B. (2019). A glucose-sensing neuron pair regulates insulin and glucagon in Drosophila. Nature 574, 559–564.

109. Jo, S., Kim, T., Iyer, V.G., and Im, W. (2008). CHARMM-GUI: a web-based graphical user interface for CHARMM. J. Comput. Chem. 29, 1859–1865.

110. Lee, J., Patel, D.S., Ståhle, J., Park, S.-J., Kern, N.R., Kim, S., Lee, J., Cheng, X., Valvano, M.A., Holst, O., et al. (2019). CHARMM-GUI Membrane Builder for complex biological membrane simulations with glycolipids and lipoglycans. J. Chem. Theory Comput. 15, 775–786.

111. Tsuyama, T., Tsubouchi, A., Usui, T., Imamura, H., and Uemura, T. (2017). Mitochondrial dysfunction induces dendritic loss via eIF2α phosphorylation. J. Cell Biol. 216, 815–834.

112. Hayashi, Y., Tsuyama, T., Usui, T., Ohbayashi, T., Kondo, T., Kokuryoh, K., Awaya, T., Katsuno, T., and Uemura, T. (2026). Roles of a Drosophila ADAM 10 transmembrane metalloprotease, Kuzbanian, in tissue architecture and function of the adult adipose tissue. Genes Cells 31, e70084.

113. Satoh, D., Sato, D., Tsuyama, T., Saito, M., Ohkura, H., Rolls, M.M., Ishikawa, F., and Uemura, T. (2008). Spatial control of branching within dendritic arbors by dynein-dependent transport of Rab5-endosomes. Nat. Cell Biol. 10, 1164–1171.

